# Glycan-Based Shaping Of The Microbiota During Primate Evolution

**DOI:** 10.1101/2021.02.10.430443

**Authors:** Sumnima Singh, Patricia Bastos-Amador, Jessica A. Thompson, Mauro Truglio, Bahtiyar Yilmaz, Silvia Cardoso, Daniel Sobral, Miguel P. Soares

**Author notes:** Equal contribution (Patricia Bastos-Amador, Jessica A. Thompson). Current address: Center for Systems Biology, Massachusetts General Hospital and Harvard Medical School, Boston, USA. Champalimaud Centre for the Unknown, Lisbon, Portugal (Patricia Bastos-Amador). Department for Biomedical Research, Inselspital, University of Bern, Bern, Switzerland (Bahtiyar Yilmaz). Universidade Nova de Lisboa, Caparica, Portugal (Daniel Sobral).

## Abstract

Genes encoding certain glycosyltransferases are thought to be under relatively high selection pressure, due to the involvement of the glycans that they synthesize in host-microbe interactions. Here we used a mouse model to investigate whether the loss of α-1,3-galactosyltransferase (*GGTA1*) function and Galα1-3Galβ1-4GlcNAcβ1-R (αGal) expression during primate evolution might have affected host-microbiota interactions. We found that *Ggta1* deletion in mice shaped the composition of the gut microbiota in relation to the bacterial species present. This occurred via an immunoglobulin (Ig)-dependent mechanism, associated with IgA targeting of αGal-expressing bacteria. Systemic infection by the Ig-shaped microbiota elicited a less severe form of sepsis than infection with the non-Ig-shaped microbiota. This suggests that in the absence of host αGal, the microbiota is shaped towards lower pathogenicity, likely providing a fitness gain to the host. We infer that high selection pressure exerted by bacterial sepsis may have contributed to increase frequency of *GGTA1* loss-of-function mutations in ancestral primates that gave rise to humans.

## Introduction

As initially proposed by J.B.S. Haldane, infectious diseases are a major driving force of natural selection (Haldane, 1949), occasionally precipitating “catastrophic-selection” events: the replacement of an entire, susceptible, parental population by mutant offspring that are resistant to a given infectious disease (Lewis, 1962). Such an event is proposed to have occurred during primate evolution between 20–30 million□years□ago, possibly in response to an airborne, enveloped virus carrying a Galα1-3Galβ1-4GlcNAc (αGal)-like glycan (Galili, 2016, 2019). If proven correct, this would contribute to our understanding of the evolutionary pressures that led to the independent fixation of several loss-of-function mutations in the *GGTA1* gene of ancestral primates (Galili et al., 1988).

Loss of the α1,3-galactosyltransferase enzyme, encoded by *GGTA1*, eliminated the expression of protein-bound αGal, allowing for immune targeting of this non-self glycan (Galili et al., 1987). This increased resistance to infection by αGal-expressing pathogens (Repik et al., 1994; Takeuchi et al., 1996), including parasites of the *Plasmodium* spp. (Soares and Yilmaz, 2016; Yilmaz et al., 2014), the causative agents of malaria, which exerted a major impact on human evolution (Allison, 1954).

We recently uncovered a fitness advantage associated with loss of *GGTA1* function in mice, which acts independently of αGal-specific immunity (Singh et al., 2020). Namely, the loss of αGal from the immunoglobulin (Ig)G-associated glycan structures increases IgG effector function (Singh et al., 2020) and resistance to bacterial sepsis, a life-threatening organ dysfunction caused by a deregulated response to infection (Singer et al., 2016), that accounts for 20% of global human mortality (Rudd et al., 2020).

The pathogenesis of sepsis is modulated by stable symbiotic associations between the host and microbial communities composed of bacteria, fungi and viruses, known as the microbiota (Rudd et al., 2020; Vincent et al., 2009). While host-microbiota interactions provide a broad range of fitness advantages to the host (Lane-Petter, 1962; Vonaesch et al., 2018), they also carry fitness costs, for example, when bacterial pathobionts (Chow et al., 2011) translocate across host epithelial barriers to promote the development of sepsis (Rudd et al., 2020; Vincent et al., 2009). On the basis of this evolutionary trade-off (Stearns and Medzhitov, 2015), the immune system may have emerged, in part, to mitigate the pathogenic effects of the microbiota (Hooper et al., 2012; McFall-Ngai, 2007). Central to this host defence strategy is the generation of copious amounts of IgA natural antibodies (NAb) targeting immunogenic bacteria in the microbiota (Macpherson et al., 2000).

IgA can undergo transepithelial secretion and target a broad but defined subset of immunogenic bacteria in the microbiota (Bunker et al., 2017; Bunker et al., 2015; Macpherson et al., 2000). In doing so, IgA can exert negative or positive selection pressure on the recognized bacteria, shaping the microbiota composition, ecology, and potentially its pathogenicity (Kubinak and Round, 2016). Negative selection can occur, for example, when IgA limit bacterial growth (Moor et al., 2017). Positive selection can occur, for example, when IgA promotes bacterial interactions with the host, favoring bacterial retention, fitness and colonization (Donaldson et al., 2018; McLoughlin et al., 2016). Moreover, IgA can interfere with cognate interactions between bacteria and resident immune cells at epithelial barriers, regulating systemic microbiota-specific immune responses characterized by the production of circulating IgM and IgG NAb (Kamada et al., 2015; Zeng et al., 2016).

Here we provide experimental evidence in mice to suggest that the fixation of *GGTA1* loss-of-function mutations during primate evolution could have exerted a major impact on the composition of their gut microbiota. In support of this notion, we found that mice in which *Ggta1* is disrupted (*Ggta1^-/-^*), mimicking human *GGTA1* loss-of-function mutations, modulate the gut microbiota composition. Shaping of the gut microbiota occurs predominantly via an Ig-dependent mechanism associated with an enhanced systemic IgA response, which, upon secretion into the gut, targets αGal-expressing bacteria in the gut microbiota. The pathogenicity of the Ig-shaped microbiota is reduced, failing to elicit the development of lethal forms of sepsis upon systemic infection. We propose that *GGTA1* loss-of-function mutations could have conferred a selective benefit during primate evolution by shaping commensal bacteria in the microbiota to mitigate the pathogenesis of sepsis.

## Results

### *Ggta1* deletion shapes the microbiota composition

We have previously established that *Ggta1^-/-^* mice harbor a distinct microbiota composition to that of wild type (*Ggta1^+/+^*) mice (Singh et al., 2020). This is illustrated by the relative abundance of specific bacterial taxa, such as an increase in Proteobacteria, Tenericutes and Verrucomicrobia as well as a reduction in Bacterioidetes and Deferribacteres phyla in *Ggta1^-/-^* mice when compared to *Ggta1^+/+^* mice (*Figure 1A, S1,2*)(Singh et al., 2020). The relative increase of Proteobacteria, a phylum containing several strains associated with pathogenic behavior, i.e. pathobionts; in the gut microbiota of *Ggta1^-/-^* mice was not, however, associated with the development of histological lesions in the gastrointestinal tract (*Figure S3A*). Absence of intestinal inflammation was assessed by the levels of fecal lipocalin-2 (Lcn-2) (*Figure S3B*) (Chassaing et al., 2012). There were also no lesions in the liver, lungs, kidney and spleen (*Figure S3C*), suggesting that *Ggta1^-/-^* mice maintain symbiotic interactions with these pathobionts, without compromising organismal homeostasis.

**Figure 1.**
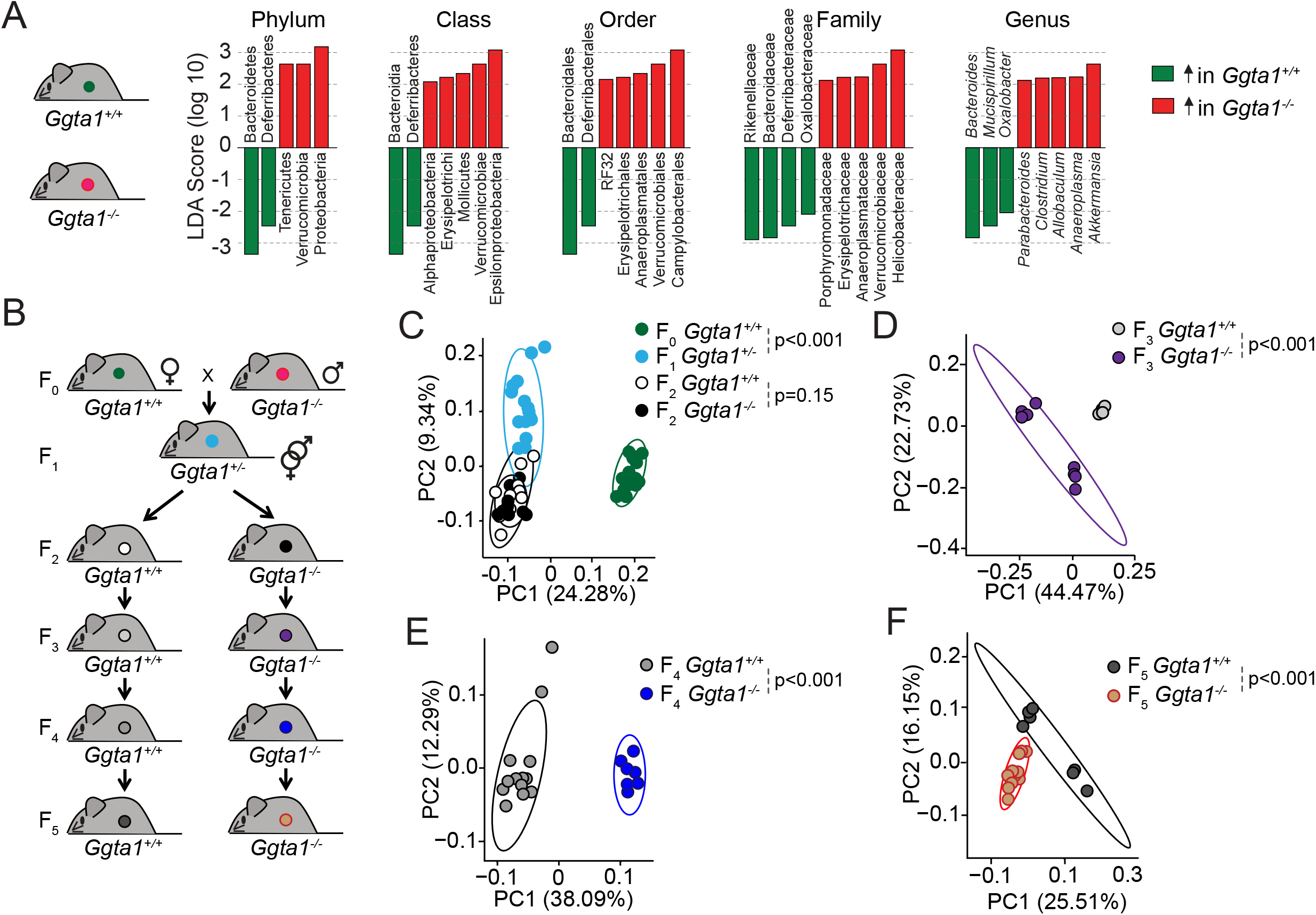
*Ggta1* deletion alters the gut microbiota. **A**) Linear discriminant analysis (LDA) scores generated from LEfSe analysis, representing taxa enriched in the fecal microbiota of *Ggta1^+/+^* (green) (n=15) and *Ggta1^-/-^* (red) (n=14) mice. **B**) Breeding strategy where F_0_ *Ggta1^-/-^* males were crossed with *Ggta1^+/+^* females to generate F_1_ *Ggta1^+/-^* mice, which were bred to generate F_2_ *Ggta1^+/+^ vs. Ggta1^-/-^* littermates. These were subsequently bred to generate F_3_ to F_5_ *Ggta1^+/+^ vs. Ggta1^-/-^* mice. Microbiota Principal Coordinate Analysis of Unweighted Unifrac distance in fecal samples from **C**) F_0_ *Ggta1^+/+^* (n=15), F_1_ *Ggta1^+/-^* (n=15), F_2_ *Ggta1^+/+^* (n=11) and F_2_ *Ggta1^-/-^* (n=10) mice, **D**) F_3_ *Ggta1^+/+^* (n=9) and *Ggta1^-/-^* (n=8) mice, **E**) F_4_ *Ggta1^+/+^* (n=13) and *Ggta1^-/-^* (n=7) mice and **F**) F_5_ *Ggta1^+/+^* (n=7) and *Ggta1^-/-^* (n=12) mice generated as in (B). Data from 1 experiment with 2-3 independent breedings/cages per genotype. Symbols (C-F) are individual mice. P values (C-F) calculated using PERMANOVA test.

To establish that the observed differences in the bacterial species present in the gut microbiota of *Ggta1^-/-^ vs. Ggta1^+/+^* mice is propelled by host genetics, we used an experimental system, whereby the microbiota is vertically transmitted over several generations (Ubeda et al., 2012) from *Ggta1^+/+^* mice to *Ggta1^-/-^* and *Ggta1^+/+^* offspring (*Figure 1B*). This approach enables the effects exerted by the host genotype on microbiota composition to predominate over those exerted by environmental factors (Gálvez et al., 2017; Vonaesch et al., 2018), diet (Sonnenburg et al., 2016), cohousing or familial transmission (Ubeda et al., 2012), albeit not accounting for putative cage effects or genetic drift (Spor et al., 2011).

The microbiota composition of F_1_ *Ggta1^+/-^* as well as F_2_ *Ggta1^+/+^* and *Ggta1^-/-^* mice diverged from that of the original F_0_ *Ggta1^+/+^* mice (*Figure 1C*). While indistinguishable in F_2_ *Ggta1^+/+^* and *Ggta1^-/-^* littermates (*Figure 1C*) (Singh et al., 2020), the microbiota composition diverged between *Ggta1^+/+^* and *Ggta1^-/-^* mice in F_3_ (*Figure 1D*), F_4_ (*Figure 1E*) and F_5_ (*Figure 1F*) generations, suggesting that the *Ggta1* genotype *per se* alters microbiota composition. There was an enrichment of some bacterial taxa, such as Helicobactereaceae, in the microbiota of F_2_ to F_5_ *Ggta1^-/-^* and *Ggta1^+/+^* mice (*Figure S4*), despite the absence of these bacteria in the original F_0_ *Ggta1^+/+^* microbiota (*Figure S4*). This suggests that while the *Ggta1* genotypes shapes the gut microbiota composition, this occurs via a process that is probably influenced by colonization by environmental pathobionts (Gálvez et al., 2017).

### *Ggta1* deletion enhances IgA responses to the gut microbiota

Considering that IgA shapes the bacterial species present in the microbiota (Bunker et al., 2015; Macpherson et al., 2018; Peterson et al., 2007), we asked whether differences in IgA responses could contribute to shape the microbiota of *Ggta1^-/-^ vs. Ggta1^+/+^* mice. In keeping with this notion, the relative levels of microbiota-reactive circulating IgA were higher in *Ggta1^-/-^ vs. Ggta1^+/+^* mice, when maintained under specific pathogen-free (SPF) but not germ-free (GF) conditions (*Figure 2A*). When maintained under SPF conditions *Ggta1^-/-^* mice had similar levels of secreted (*Figure S5A*) and circulating (*Figure S5B*) total IgA, when compared to *Ggta1^+/+^* mice. These were reduced to a similar extent under GF conditions (*Figure S5A-B*). This suggests that *Ggta1* deletion preferentially enhances the microbiota-specific IgA response without interfering with total IgA.

**Figure 2.**
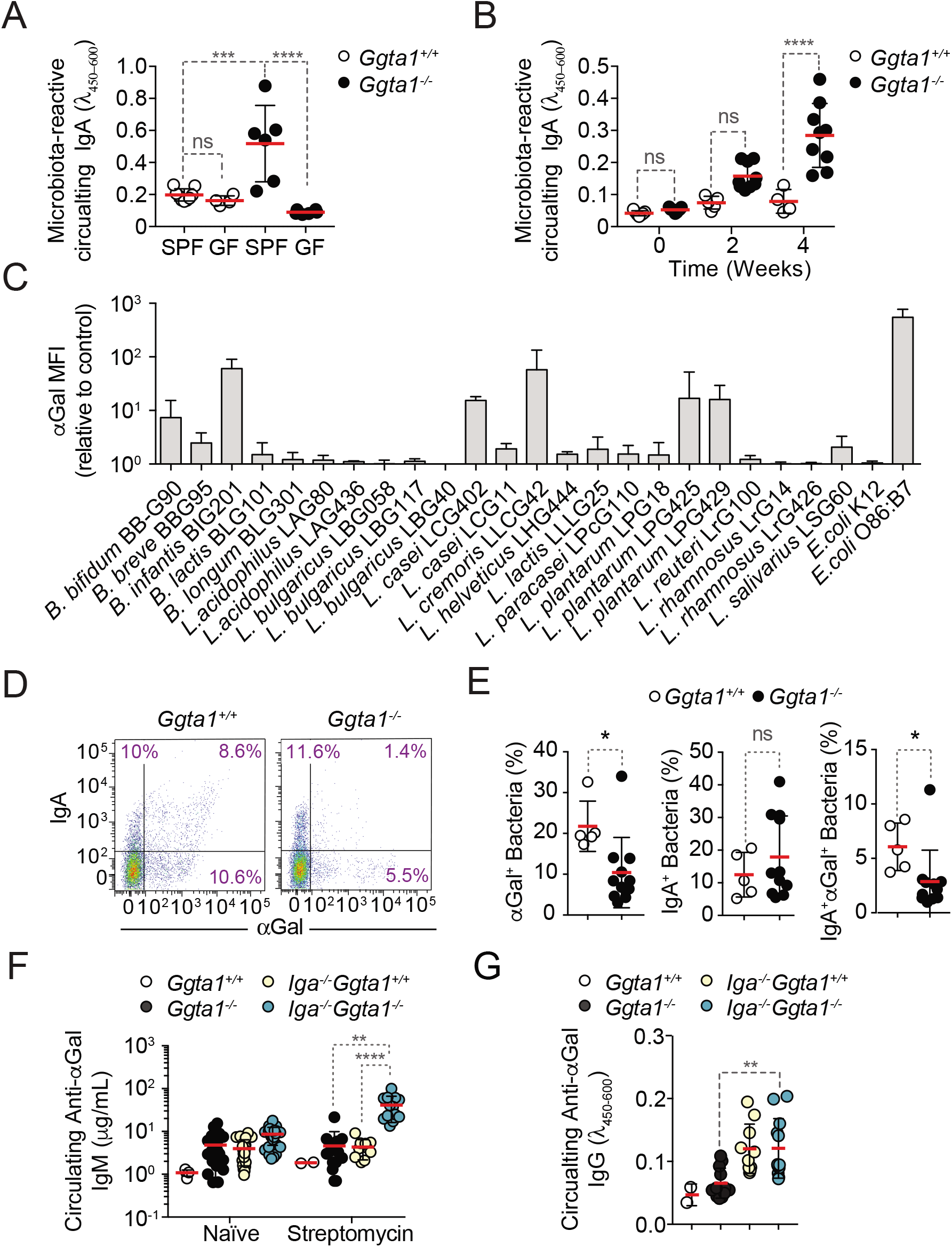
*Ggta1* deletion enhances IgA responses to the gut microbiota. **A**) Relative binding of IgA in the serum of *Ggta1^+/+^* (n=10) and GF *Ggta1^+/+^* (n=5) mice to fecal extract from *Ggta1^+/+^* mice, and *Ggta1^-/-^* (n=6) and GF *Ggta1^-/-^* (n=6) mice to fecal extract from *Ggta1^-/-^* mice; 1 experiment. **B**) Relative binding of IgA in the serum of GF *Ggta1^+/+^* (n=7) and GF *Ggta1^-/-^* (n=10) mice to cecal extract from *Ggta1^-/-^* mice at indicated time-points after colonization with cecal extract from *Ggta1^-/-^* mice; 2 experiments. **C**) Median Fluorescence Intensity (MFI) of αGal^+^ bacterial strains stained with BSI-B4 lectin relative to unstained control; 7 experiments. **D**) Representative flow cytometry plots showing bacteria stained for IgA and αGal in the small intestinal content of *Ggta1^+/+^* (n=5) and *Ggta1^-/-^* (n=11) mice; 4 independent experiments. **E**) Quantification of αGal^+^, IgA^+^ and IgA^+^αGal^+^ bacteria in the same samples as in (D). **F**) Concentration of anti-αGal IgM in serum of *Ggta1^+/+^* (n=2), *Ggta1^-/-^* (n=12), *Iga^-/-^Ggta1^+/+^* (n=10) and *Iga^-/-^Ggta1^-/-^* (n=12) mice before and after streptomycin treatment, 2 experiments. **G**) Concentration of anti-αGal IgG, in the same mice as (F). Symbols (A, B, E, F, G) are individual mice. Red bars (A, B, E, F, G) correspond to mean values. Error bars (A, B, C, E, F, G) correspond to SD. P values in (A, B, F, G) calculated using Kruskal-Wallis test using Dunn’s multiple comparisons test and in (E) using Mann-Whitney test. *P < 0.05, **P < 0.01, ***P < 0.001, ****P < 0.0001, ns: not significant.

In further support of the notion that *Ggta1* deletion modulates the production microbiota-specific IgA, colonization of GF *Ggta1^-/-^* mice with the microbiota from *Ggta1^-/-^* mice elicited the production of higher levels of microbiota-reactive circulating IgA, compared to GF *Ggta1^+/+^* mice colonized by the same microbiota (*Figure 2B*). This difference was not observed upon colonization of GF *Ggta1^-/-^ vs. Ggta1^+/+^* mice by a microbiota isolated from *Ggta1^+/+^* mice (*Figure S5C*). This suggests that the enhanced microbiota-reactive IgA response of *Ggta1^-/-^ vs. Ggta1^+/+^* mice is induced by immunogenic bacteria present specifically in the microbiota of *Ggta1^-/-^* mice.

Of note, the levels of circulating IgA were reduced in *Tcrβ^-/-^Ggta1^-/-^* mice lacking T cells, when compared to *Ggta1^-/-^* mice (*Figure S5D*), suggesting that the production of circulating IgA NAb in *Ggta1^-/-^* mice occurs, in part, via a T-cell dependent mechanism, which is consistent with previous reports in *Ggta1^+/+^* mice (Bunker et al., 2017; Fagarasan et al., 2010; Macpherson et al., 2000).

We then asked whether *Ggta1^-/-^* mice shape their microbiota via a mechanism associated with immune targeting of αGal-expressing bacteria. Consistent with a number of bacteria in the human gut microbiota carrying genes orthologous to the mammalian α1,3-galactosyltransferase (Montassier et al., 2019), several human probiotic bacteria expressed αGal-like glycans at the cell surface when cultured *in vitro (Figure 2C, S6*). Subsequent *in vivo* analyses demonstrated that approximately 20% of the bacteria in the small intestine of *Ggta1^+/+^* mice expressed αGal-like glycans at the cell surface (*Figure 2D,E, S5E*). Nearly 30% of these αGal^+^ bacteria were immunogenic (*Figure 2D,E*), as defined by the detection of surface-bound IgA (Palm et al., 2014), which predominantly targets bacteria in the small intestine (Bunker et al., 2017; Bunker et al., 2015). These IgA^+^αGal^+^ bacteria accounted for roughly 50% of all the immunogenic (IgA^+^) bacteria in the small intestine (*Figure 2D,E*). These were also present, although at a lower extent, in the cecum, colon and feces (*Figure S5F*).

*Ggta1^-/-^* mice harbored a relatively lower percentage of immunogenic IgA^+^αGal^+^ bacteria in the small intestine, when compared to *Ggta1^+/+^* mice (*Figure 2D,E*), while the percentage of immunogenic IgA^+^ bacteria was similar in *Ggta1^-/-^ vs. Ggta1^+/+^* mice (*Figure 2D,E*). This is consistent with the idea of a specific mechanism altering the microbiota of *Ggta1^-/-^* mice that presumably, at least in part, involves targeting immunogenic αGal^+^ bacteria by IgA. Whether this mechanism involves the recognition of bacterial αGal-like glycans by IgA is not clear.

We than compared the effect of IgA on the levels of systemic IgM and/or IgG NAb directed against antigens expressed by bacteria present in the microbiota (Kamada et al., 2015; Zeng et al., 2016). Induction of dysbiosis by streptomycin, increased the levels of circulating αGal-specific IgM and IgG in *Iga^-/-^Ggta1^-/-^ vs. Iga^+/+^Ggta1^-/-^* mice (*Figure 2F,G*). This illustrates again that the IgA response of *Ggta1^-/-^* mice is distinct from that of *Ggta1^+/+^* mice, reducing systemic IgM and IgG responses against antigens expressed by bacteria in the microbiota, as illustrated for αGal-like glycans. Presumably this occurs via a mechanism, whereby IgA prevents αGal^+^ bacteria or bacterial products associated to αGal from translocating across epithelial barriers and inducing systemic immune responses against this glycan.

### *Ggta1* deletion shapes the gut microbiota via an antibody (Ig)-dependent mechanism

Maintenance of microbiota composition across generations is sustained via maternal IgG transfer to the offspring through the placenta during the fetal period, and maternal IgM, IgG and IgA transfer via lactation during the neonatal period (Gensollen et al., 2016; Koch et al., 2016). Over time, newborns generate IgA that target immunogenic bacteria in the microbiota (McLoughlin et al., 2016; Moor et al., 2017), shaping its composition throughout adult life (Kawamoto et al., 2014). We hypothesized that the mechanism via which *Ggta1* deletion shapes the gut microbiota involves targeting of immunogenic bacteria by both maternally-and offspring-derived immunoglobulins (Ig). To test this hypothesis, we used a similar experimental approach to that described above (*Figure 1B*) (Ubeda et al., 2012), whereby the microbiota from *Ggta1^+/+^* mice was vertically transmitted to *Ggta1^+/+^* or *Ggta1^-/-^* offspring that express Ig (*J_h_t^+/+^*) or not (*J_h_t^-/-^) (Figure 3A*). Crossing of F_0_ *J_h_t^+/+^Ggta1^+/+^* (female) with *J_h_t^-/-^Ggta1^-/-^* (male) mice produced F_1_ *J_h_t^+/-^Ggta1^+/-^* mice, which were interbred to generate F_2_ *J_h_t^+/+^Ggta1^+/+^, J_h_t^+/+^Ggta1^-/-^, J_h_t^-/-^Ggta1^+/+^* and *J_h_t^-/-^ Ggta1^-/-^* mice, harboring a microbiota composition indistinguishable among genotypes (*Figure S7A,B*). Consistent with our previous observations (*Figure 1C*) (Singh et al., 2020), this suggests that maternal Ig transfer predominates over offspring Ig production in shaping the offspring microbiota, regardless of the genotype (Ubeda et al., 2012).

**Figure 3.**
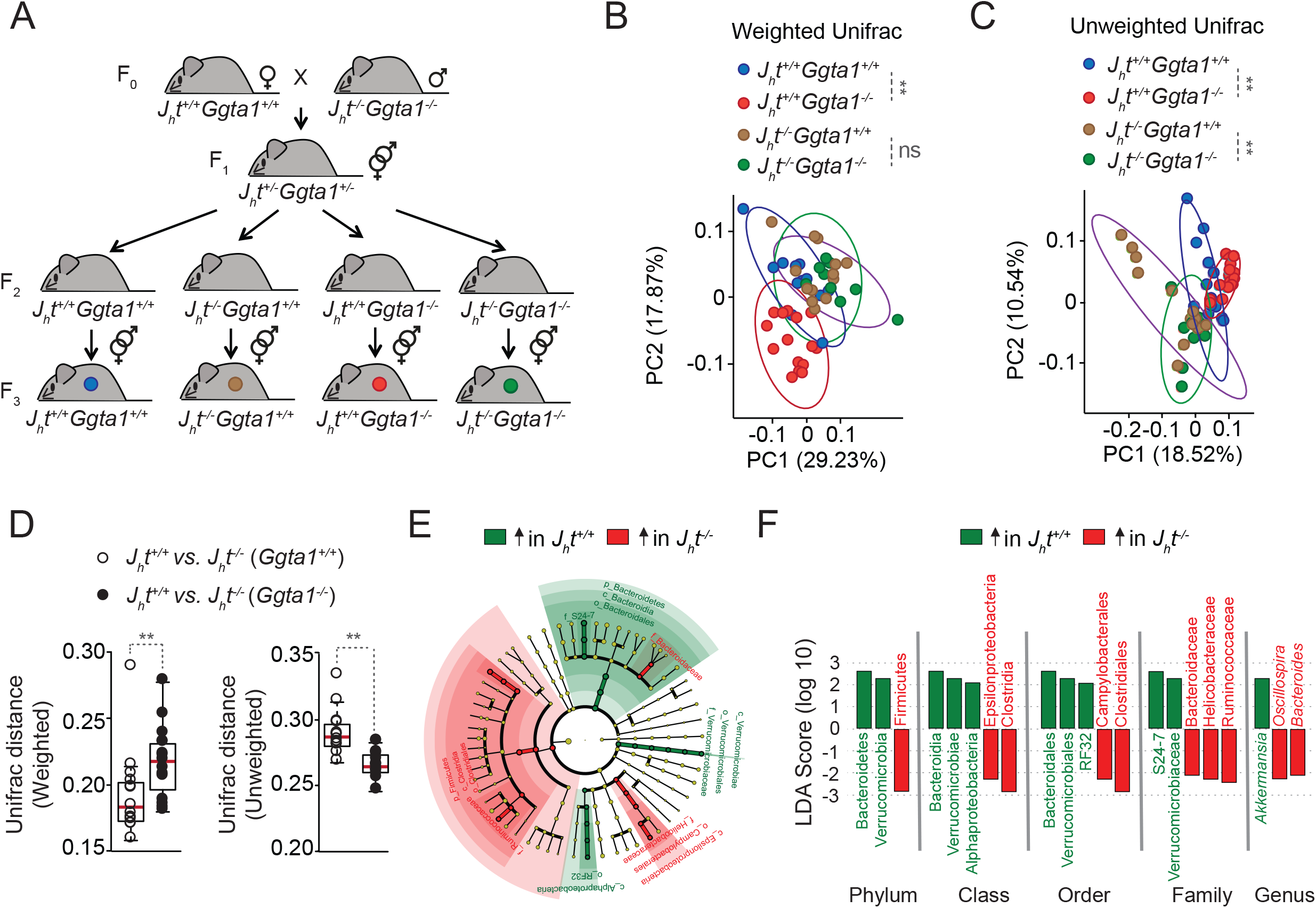
*Ggta1* deletion shapes the gut microbiota via an Ig-dependent mechanism. **A**) Breeding strategy where F_0_ *J_h_t^-/-^Ggta1^-/-^* males were crossed with *J_h_t^+/+^Ggta1^+/+^* females to generate F_1_ *J_h_t^+/-^Ggta1^+/-^* mice, which were bred to generate F_2_ and F_3_ *J_h_t^+/+^Ggta1^+/+^, J_h_f^-/-^Ggta1^+/+^, J_h_t^+/+^Ggta1^-/-^ and J_h_t^-/-^Ggta1^-/-^* mice. Microbiota Principal Coordinate Analysis of **B**) Weighted and **C**) Unweighted Unifrac and **D**) Distance of Weighted and Unweighted Unifrac of 16S rRNA amplicons, in fecal samples from F_3_ *J_h_t^+/+^Ggta1^+/+^* (n=13) *vs. J_h_t^-/-^Ggta1^+/+^* (n=16) mice and F_3_ *J_h_t^+/+^Ggta1^-/-^* (n=13) *vs. J_h_t^-/-^Ggta1^-/-^* (n=11) mice generated as in (A). **E**) Cladogram and **F**) Linear discriminant analysis (LDA) scores generated from LEfSe analysis, representing taxa enriched in the fecal microbiota of the same mice as in (B-D). Data from 1 experiment with 2-3 independent breedings/cages per genotype. Symbols (B, C, D) are individual mice. Red bars (D) correspond to mean values. Error bars (D) correspond to SD. P values in (B, C) calculated using PERMANOVA and in (D) using Mann-Whitney test. **P < 0.01, ns: not significant.

To dissect the relative contribution of the *Ggta1* from the *Ig* genotype in shaping the microbiota, F_2_ littermates were interbred to generate F_3_ offspring carrying a microbiota targeted by antibodies (*J_h_t^+/+^Ggta1^+/+^* and *J_h_t^+/+^Ggta1^-/-^*) or not (*J_h_t^-/-^Ggta1^+/+^* and *J_h_t^-/-^Ggta1^-/-^) (Figure 3A*). Consistent with our previous observations (*Figure 1*), there was a marked separation of the microbiota community structure between F_3_ *J_h_t^+/+^Ggta1^-/-^ vs. J_h_t^+/+^Ggta1^+/+^* mice, as assessed by Principal Coordinate Analyses (PCA) for Weighted and Unweighted Unifrac (*Figure 3B,C*). Considering that Weighted Unifrac accounts for the relative abundance of the bacterial taxa while Unweighted Unifrac accounts for only the presence or absence of the taxa in the microbial community (Lozupone and Knight, 2005), they represent differences exerted by high and low abundant bacterial taxa respectively. This suggests that *Ggta1* deletion shapes the microbiota when maternal and/or offspring-derived antibodies (*i.e*., Ig) are present.

LefSe analysis showed that the microbiota from *J_h_t^+/+^Ggta1^-/-^* mice was enriched in specific bacterial taxa, including Proteobacteria, as compared to the microbiota from *J_h_t^+/+^Ggta1^+/+^* mice (*Figure S7C*). This suggests that *Ggta1* deletion favors gut colonization by pathobionts in an Ig-dependent manner.

We then asked whether Ig are functionally involved in shaping the microbiota composition of *Ggta1^-/-^ vs. Ggta1^+/+^* mice. In the absence of Ig (*J_h_t^-/-^*), there were no differences in the microbiota composition of F_3_ *J_h_t^-/-^Ggta1^-/-^ vs. J_h_t^-/-^Ggta1^+/+^* mice, as assessed by PCA for Weighted Unifrac (*Figure 3B*). This suggests that shaping of highly abundant taxa in the microbiota of *Ggta1^-/-^ vs. Ggta1^+/+^* mice occurs via an Ig-dependent mechanism. In contrast, the microbiota composition of F_3_ *J_h_t^-/-^Ggta1^-/-^* mice remained distinct from that of *J_h_t^-/-^Ggta1^+/+^* mice, as assessed by PCA for Unweighted Unifrac (*Figure 3C*). This suggests that shaping of low abundance bacterial taxa in the microbiota of *Ggta1^-/-^ vs. Ggta1^+/+^* mice occurs in an Ig-independent manner.

To address the extent to which *Ggta1* deletion promotes shaping of the microbiota via an Ig-dependent mechanism, we compared the Unifrac distances between *J_h_t^+/+^Ggta1^+/+^* and *J_h_t^-/-^Ggta1^+/+^* mice *vs*. *J_h_t^+/+^Ggta1^-/-^* and *J_h_^-/-^Ggta1^-/-^* mice, similar to what was previously described (Lozupone and Knight, 2005). The Weighted Unifrac distance of microbiota from *J_h_t^+/+^Ggta1^-/-^ vs. J_h_t^-/-^Ggta1^-/-^* mice was higher than that from *J_h_t^+/+^Ggta1^+/+^ vs. J_h_t^-/-^Ggta1^+/+^* mice (*Figure 3D*). This suggests that the relative impact of Ig on the gut microbiota community structure exerted by highly abundant bacterial taxa is enhanced in *Ggta1^-/-^ vs. Ggta1^+/+^* mice. This was confirmed by LefSe analysis showing an enhanced Ig-dependent enrichment of several bacterial taxa in the microbiota of *Ggta1^-/-^* mice (*Figure 3E-F*), compared to that of *Ggta1^+/+^* mice (*Figure S7D*). In contrast, the Unweighted Unifrac distance of *J_h_t^+/+^Ggta1^+/+^ vs. J_h_t^-/-^Ggta1^+/+^* mice was higher compared to *J_h_t^+/+^Ggta1^-/-^ vs. J_h_t^-/-^ Ggta1^-/-^* mice (*Figure 3D*). This suggests that the relative impact of Ig on the microbiota community structure of low abundant bacterial taxa is less pronounced for *Ggta1^-/-^ vs. Ggta1^+/+^* mice.

We then asked whether Ig exert a higher impact on the relative abundance of pathobionts in the gut microbiota of *Ggta1^-/-^ vs. Ggta1^+/+^* mice. In strong support of this notion, the gut microbiota from *J_h_t^-/-^Ggta1^-/-^* mice, lacking Ig, was enriched with Helicobactereaceae family from Proteobacteria phylum as compared to *J_h_t^+/+^Ggta1^-/-^* mice expressing Ig (*Figure 3E, F*). This is consistent with our previous finding that the gut microbiota of *Rag2^-/-^Ggta1^-/-^* mice, lacking adaptive immunity, is highly enriched in Proteobacteria, including *Helicobacter* (Singh et al., 2020). Of note, this was not observed in *J_h_t^-/-^Ggta1^+/+^ vs. J_h_t^+/+^Ggta1^+/+^* mice (*Figure S7D*). These data suggest that the absence of host αGal favors colonization of the gut microbiota by pathobionts, the expansion of which is restrained by Ig.

### *Ggta1* deletion reduces microbiota pathogenicity

We then asked whether the Ig-dependent shaping of the microbiota in *Ggta1^-/-^* mice affects the pathogenesis of sepsis due to systemic infections emanating from gut microbes. *Ggta1^-/-^* mice were protected against systemic infections (i.p.) by a cecal inoculum isolated from *Rag2^-/-^Ggta1^-/-^* mice, reflecting a microbiota not shaped by Ig (*Figure S8A,B*). This is consistent with our previous finding showing *Ggta1* deletion enhances protection against systemic bacterial infections (i.p.) via a mechanism involving IgG NAb (Singh et al., 2020). Moreover, *Ggta1^-/-^* mice were also protected against a systemic infection (i.p.) by a cecal inoculum isolated from *Ggta1^-/-^* mice, reflecting their own Ig-shaped microbiota (*Figure S8A,B*). This is in keeping with the previously shown enhanced protection of *Ggta1^-/-^* mice against cecal ligation and puncture (Singh et al., 2020). Surprisingly *J_h_t^-/-^Ggta1^-/-^* mice lacking B cells (*Figure S8C), Tcrβ^-/-^ Ggta1^-/-^* mice lacking α/β T cells (*Figure S8D*) and *Rag2^-/-^Ggta1^-/-^* mice lacking B and T cells (*Figure S8E*) remained protected against systemic infections by the cecal inoculum isolated from *Ggta1^-/-^* mice. This suggests that the previously described IgG-dependent mechanism that protects *Ggta1^-/-^* mice from a systemic infection by a cecal inoculum isolated from *Rag2^-/-^Ggta1^-/-^* mice (Singh et al., 2020) is distinct from that protecting *Ggta1^-/-^* mice from a systemic infection by their own cecal inoculum. We reasoned that this can be explained by a reduction of the overall pathogenicity of the Ig-shaped microbiota of *Ggta1^-/-^* mice, compared to the non-Ig-shaped microbiota of *Rag2^-/-^Ggta1^-/-^* mice. To test this hypothesis, we compared the lethal outcome of *Rag2^-/-^Ggta1^-/-^* mice upon a systemic infection by Ig-shaped *vs*. a non-Ig-shaped microbiota.

*Rag2^-/-^Ggta1^-/-^* mice remained protected against systemic infections by the cecal inoculum isolated from *Ggta1^-/-^* mice, while succumbing to a systemic infection by a cecal inoculum isolated from *Rag2^-/-^Ggta1^-/-^* mice (*Figure 4A,B*). Lethality was associated with a 10^6^-10^7^-fold increase in bacterial load (*Figure 4C*), suggesting that the Ig-shaped microbiota of *Ggta1^-/-^* mice is less pathogenic, when compared to the non-Ig-shaped microbiota from *Rag2^-/-^Ggta1^-/-^* mice.

**Figure 4.**
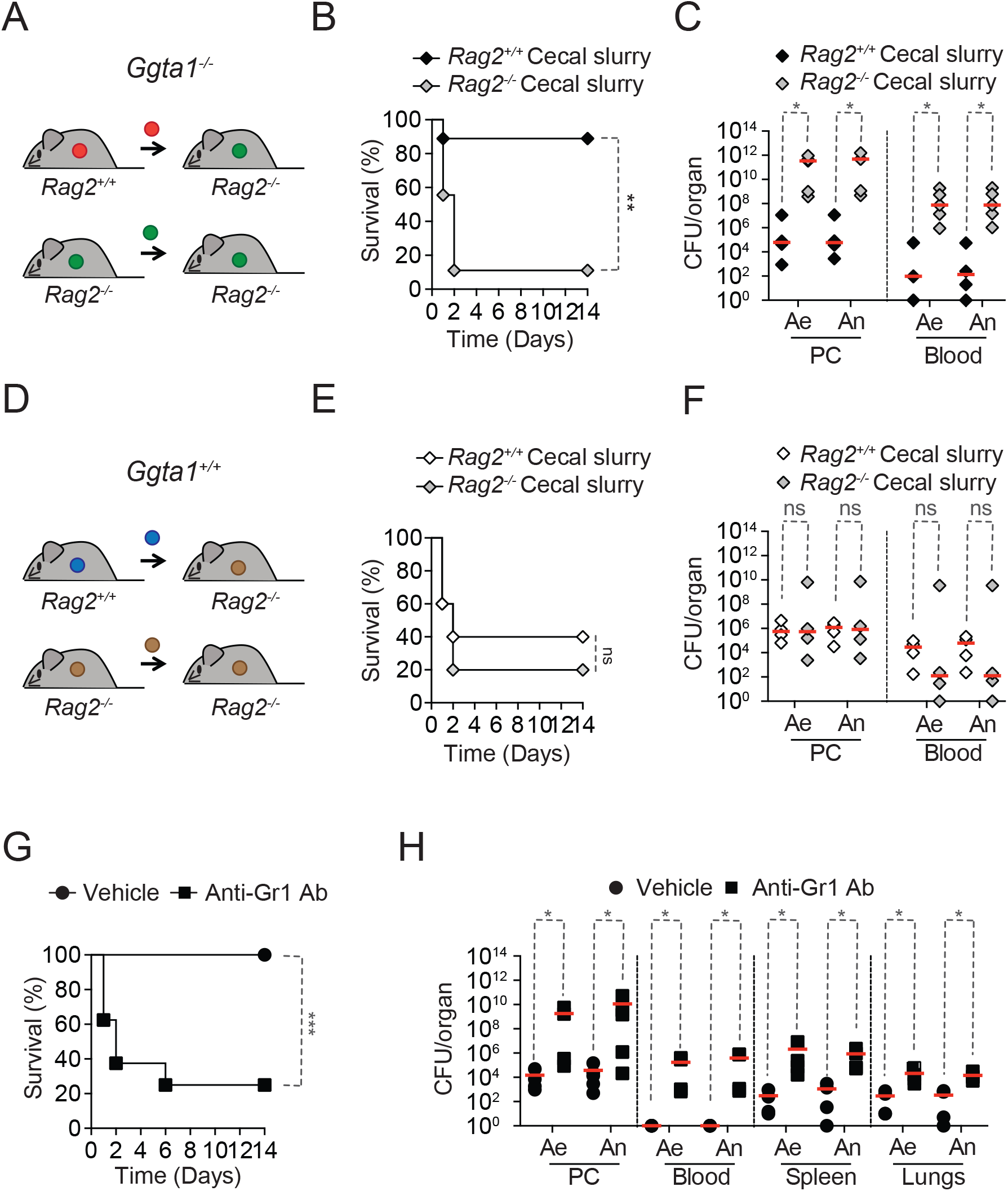
*Ggta1* deletion reduces microbiota pathogenicity. **A**) Schematic showing infection of *Rag2^-/-^Ggta1^-/-^* mice with a cecal inoculum from either *Ggta1^-/-^* or *Rag2^-/-^Ggta1^-/-^* mice. **B**) Survival of *Rag2^-/-^Ggta1^-/-^* (n=9) mice after infection with a cecal inoculum from *Ggta1^-/-^* mice, and of *Rag2^-/-^Ggta1^-/-^* (n=9) mice after infection with a cecal inoculum from *Rag2^-/-^Ggta1^-/-^* mice; 2 experiments. **C**) Colony forming units (CFU) of aerobic (Ae) and anaerobic (An) bacteria in *Rag2^-/-^Ggta1^-/-^* mice (n=5 per group), 24 hours after infection as in (B); 2 experiments. **D**) Schematic showing infection of *Rag2^-/-^Ggta1^+/+^* mice with a cecal inoculum from either *Ggta1^+/+^* or *Rag2^-/-^ Ggta1^+/+^* mice. **E**) Survival of *Rag2^-/-^Ggta1^+/+^* (n=5) mice after infection with a cecal inoculum from *Ggta1^+/+^* mice, and of *Rag2^-/-^ Ggta1^+/+^* (n=5) mice after infection with a cecal inoculum from *Rag2^-/-^Ggta1^+/+^* mice; 1 experiment. **F**) CFU of Ae and An bacteria in *Rag2^-/-^Ggta1^-/-^* mice (n=4 per group), 24 hours after infection as in (E); 1 experiment. **G**) Survival of *Ggta1^-/-^* mice receiving vehicle (PBS) (n=9) or Anti-Gr1 Ab (n=8), 24 hours before infection with cecal inoculum from *Ggta1^-/-^* mice; 2 experiments. **H**) CFU of Ae and An bacteria in *Ggta1^-/-^* mice receiving vehicle (PBS) (n=4-5) or Anti-Gr1 Ab (n=4-5), 24 hours after infection as in (G); 5 experiments. Symbols (C, F, H) represent individual mice. Red lines (C, F, H) correspond to median values. P values in (B, E, G) calculated with log-rank test and in (C, F, H) with Mann-Whitney test. Peritoneal cavity (PC). *P < 0.05, **P < 0.01, ***P < 0.001, ns: not significant.

We then asked whether the reduction of microbiota pathogenicity imposed by the adaptive immune system of *Ggta1^-/-^* mice is also operational in *Ggta1^+/+^* mice. However, *Rag2^-/-^Ggta1^+/+^* mice succumbed to the same extent to systemic infection (i.p.) with a cecal inoculum isolated from either *Rag2^+/+^Ggta1^+/+^ vs. Rag2^-/-^Ggta1^+/+^* mice (*Figure 4D-E*), developing similar bacterial loads (*Figure 4F*). This suggests that the mechanism via which the adaptive immune system of *Ggta1^-/-^* mice shapes and reduces the pathogenicity of its microbiota is not operational in *Ggta1^+/+^* mice.

Having established that in the absence of adaptive immunity, *Ggta1^-/-^* mice are resistant against systemic infection by a low pathogenic Ig-shaped microbiota (*Figure S8C-E*), we asked whether the mechanism of resistance relies on the innate immune system. Depletion of Ly6C^+^/Ly6G^+^ myeloid cells (i.e., polymorphonuclear cells and inflammatory monocytes) using an anti-GR1 monoclonal Ab (*Figure S8F*) (Daley et al., 2008) increased the susceptibility of *Ggta1^-/-^* mice to systemic infection by their cecal inoculum (*Figure 4G*). This was associated with a 10^2^-10^5^-fold increase in bacterial load, compared to control *Ggta1^-/-^* mice (*Figure 4H*). Of note, monocyte/macrophage depletion by Clodronate liposomes (*Figure S8G*) (Sunderkötter et al., 2004a; Sunderkötter et al., 2004b) failed to compromise the survival of *Ggta1^-/-^* mice upon infection by the same cecal inoculum (*Figure S8H*). This suggests that polymorphonuclear cells are essential for resistance against systemic infection emanating from the less pathogenic Ig-shaped microbiota of *Ggta1^-/-^* mice.

## Discussion

While loss-of-function mutations in genes encoding glycosyltransferases can provide major fitness advantages against infection, these can also compromise the physiologic functions of the eliminated self-glycan, as illustrated by the occurrence of reproductive senescence upon the loss of *GGTA1* function in mice (Singh et al., 2020). In an evolutionary context, such a trade-off could explain why these loss-of-function mutations are rare, and in some cases unique, to the human lineage, as illustrated for the loss of CMP-*N*-acetylneuraminic acid hydroxylase (*CMAH*) function, which eliminated sialic acid *N*-glycolylneuraminic acid (Neu5Gc) expression in humans (Ghaderi et al., 2011).

Ancestral hominids co-evolved with commensal bacteria in their microbiota over millions of years (Dethlefsen et al., 2007; Huttenhower et al., 2012; Moeller et al., 2016). The fitness advantages associated with these stable symbiotic associations include, among others, an overall optimization of nutrient intake from diet, regulation of different aspects of organismal metabolism, and colonization resistance against pathogenic bacteria (Buffie and Pamer, 2013). Here we propose that the loss of αGal expression, as it occurred during primate evolution, might have exerted a major impact on the nature of the symbiotic associations with bacteria in the microbiota. In support of this notion, *Ggta1* deletion in mice was associated with major changes in the composition of the gut microbiota, in relation to the bacterial species present, over several generations (*Figure 1*). This occurred under experimental conditions of exposure to environmental pathobionts while minimizing the relative impact exerted by other environmental factors on microbiota composition (Ubeda et al., 2012), arguing that the *Ggta1* genotype modulates the microbiota composition.

The mechanism via which *Ggta1* deletion shapes the bacterial species present the gut microbiota is associated with targeting of αGal-expressing bacteria by IgA (*Figure 2*). Consistent with this notion, a relatively large proportion of probiotic bacteria as well as bacteria present in the mouse microbiota express αGal-like glycans at the cell surface (*Figure 2*). About 30% of these carry IgA at the cell surface in the mouse microbiota and, as such, are considered as immunogenic (Palm et al., 2014). The proportion of IgA^+^αGal^+^ bacteria was reduced in the microbiota of *Ggta1*-deleted mice and was presumably eliminated (*Figure 2*). This suggests that *Ggta1* deletion probably broadens bacterial recognition by IgA to include immunogenic αGal^+^ bacteria, which as a result are probably purged from the microbiota. Whether this occurs via a mechanism involving the recognition of αGal-like glycans and/or other related epitopes expressed at the surface of these bacteria by IgA has not been established. Of note, these are not mutually exclusive possibilities since: i) IgA can target αGal-like glycans and modulate bacterial pathogenicity (Hamadeh et al., 1995a; Hamadeh et al., 1995b), ii) IgA are poly-reactive and can target common antigens expressed by bacteria (Bunker et al., 2017; Bunker et al., 2015) and iii) αGal-reactive antibodies present a degree of poly-reactivity against non-αGal related bacterial epitopes (Bernth Jensen et al., 2020).

The identity of the αGal^+^ bacteria targeted by IgA remains elusive but likely includes Gram-negative pathobionts from the *Enterobacteriaceae* family, as demonstrated for *Escherichia* (*E*.) *coli* O86:B7 (Yilmaz et al., 2014), which expresses the αGal-like glycan Galα1-3Gal(Fucα1-2)β1-3GlcNAcβ1-4Glc as part of the lipopolysaccharide (LPS) O-antigen (Guo et al., 2005). Of note, this pathobiont can induce a systemic αGal-specific NAb response in humans (Springer and Horton, 1969) as well as in *Ggta1*-deleted mice, which is protective against infection by pathogens expressing αGal-like glycans (Yilmaz et al., 2014). The finding that several commensal bacteria in the human gut microbiota express αGal-like glycans (*Figure 2, S5*) suggests that other bacteria might contribute to this protective response.

Our findings also suggest that *Ggta1* deletion shapes the bacterial community structure among highly abundant bacterial taxa in the microbiota via an Ig-dependent mechanism, and among low abundant bacterial taxa independently of Ig (*Figure 3*). Presumably, the shaping of highly abundant taxa by Ig occurs via a mechanism that involves not only IgA produced by the offspring, but also maternal IgG transferred through the placenta as well as IgM, IgG and IgA transferred through maternal milk (Koch et al., 2016). These regulate neonatal innate (Gomez de Agüero et al., 2016) and adaptive (Koch et al., 2016) immunity, shaping the offspring microbiota composition (Gensollen et al., 2016; Koch et al., 2016). Whether this occurs via targeting of αGal-like glycans, as discussed above, and/or via other bacterial antigens expressed by immunogenic bacteria has not been established.

Shaping of lowly abundant bacterial taxa independently of Ig (*Figure 3*) suggests that other mechanisms contribute to shape the microbiota of *Ggta1*-deleted mice. These probably include the modulation of nutritional or spatial niches due to the loss of αGal from complex glycosylated structures present at epithelial barriers, such as the mucus, as demonstrated for other glycans (Pickard et al., 2014).

The selective pressure exerted by the adaptive immune system of *Ggta1^-/-^* mice on the bacterial species present in the microbiota probably allows for the establishment of a more diverse ecosystem containing pathobionts (Singh et al., 2020), such as Helicobactereaceae (*Figure 1, S1,2*). These can elicit the production of antibodies targeting and exerting negative selective pressure over other bacterial symbionts and releasing ecological niches, thus further shaping the microbiota. Expansion of these pathobionts in the microbiota of *Ggta1^-/-^* mice is restrained by Ig, which probably explains the lack of associated pathology (*Figure S3*). This suggests that the loss of *GGTA1* function in ancestral primates fostered mutualistic interactions with more diverse bacterial ecosystems, incorporating pathobionts such as *Helicobacter pylori*, associated with fitness costs (Atherton and Blaser, 2009) and gains (Linz et al., 2007).

Our findings suggest that the Ig-shaping of the bacterial species present in the gut microbiota of *Ggta1^-/-^* mice reduces its pathogenicity, as illustrated by the lethal outcome of systemic infections by the Ig-shaped *vs*. non-shaped microbiota (*Figure 4A-F, S8A,B*). This reduction in pathogenicity makes that the effector mechanism underlying resistance against systemic infection by the Ig-shaped microbiota does not rely on the adaptive immune system (*Figure S8C-E*), but instead on cellular components of the innate immune systems, namely, neutrophils (*Figure 4G,H*).

The notion of host mechanisms shaping the microbiota composition towards a reduction of its pathogenicity is in keeping with host microbial interactions not being hardwired but instead shifting between symbiotic to pathogenic depending on host and microbe cooperative behaviors (Ayres, 2016; Vonaesch et al., 2018). For example, when restricted to the microbiota, bacterial pathobionts can behave as commensals posing no pathogenic threat to the host (Vonaesch et al., 2018), while triggering sepsis (Haak and Wiersinga, 2017) upon translocation across epithelial barriers (Caruso et al., 2020). The high fitness cost imposed to modern humans by sepsis (Rudd et al., 2020) suggests that mutations shaping the composition of the microbiota towards a reduced capacity to elicit sepsis should be associated with major fitness advantages. Our findings suggest that loss-of-function mutations in *GGTA1* act in such a manner.

In conclusion, protective immunity against αGal-expressing pathogens was likely a major driving force in the natural selective events that led to the fixation of loss-of-function mutations in the *GGTA1* gene of ancestral Old World primates (Galili, 2016; Soares and Yilmaz, 2016). Moreover, in the absence of αGal, the glycan structures associated with the Fc portion of IgG, can increase IgG-effector function and resistance to bacterial infections, irrespectively of αGal-specific immunity (Singh et al., 2020). The net survival advantage against infection provided by these traits came alongside the emergence of reproductive senescence (Singh et al., 2020), a major fitness cost presumably outweighed by endemic exposure to highly virulent pathogens (Singh et al., 2020). Here we provide experimental evidence for yet another fitness advantage against infection, associated with the loss of *GGTA1*, driven by Ig-shaping and reduction of microbiota pathogenicity. We infer that ancestral Old World primates carrying loss-of-function mutations in *GGTA1* probably shaped their microbiota to minimize its pathogenic effect, providing a major fitness advantage against sepsis.

## Materials and Methods

### Mice

Mice were used in accordance with protocols approved by the Ethics Committee of the Instituto Gulbenkian de Ciência (IGC) and Direção Geral de Alimentação e Veterinária (DGAV), following the Portuguese (Decreto-Lei no. 113/2013) and European (directive 2010/63/EU) legislation for animal housing, husbandry and welfare. C57BL/6J wild-type (*Ggta1^+/+^), Ggta1^-/-^* (Tearle et al., 1996), *J_h_t^-/-^Ggta1^-/-^* (Gu et al., 1993)*, Tcrβ^-/-^Ggta1^-/-^* (Yilmaz et al., 2014), *μs^-/-^Ggta1^-/-^*(Yilmaz et al., 2014), *Iga^-/-^Ggta1^-/-^* (Singh et al., 2020) and *Rag2^-/-^Ggta1^-/-^*(Singh et al., 2020) mice were used. Mice were bred and maintained under specific pathogen-free (SPF) conditions (12 h day/night, fed *ad libitum*), as described (Yilmaz et al., 2014). Germ-free (GF) C57BL/6J *Ggta1^+/+^* and *Ggta1^-/-^* animals were bred and raised in the IGC gnotobiology facility in axenic isolators (La Calhene/ORM), as described (Yilmaz et al., 2014) (Singh et al., 2020). Sterility of food, water, bedding, oral swab and feces were confirmed as described (Singh et al., 2020). Both male and female mice were used for all experiments. All animals were studied between 9-16 weeks of age unless otherwise indicated.

### Breeding experiments

Vertical transmission of the microbiota from *Ggta1^+/+^* mice to *Ggta1^-/-^* and *Ggta1^+/+^* offspring over several generations was achieved, as described (Ubeda et al., 2012) (Singh et al., 2020). Briefly, two or more breeding pairs were established, consisting of two *Ggta1^+/+^* females and one *Ggta1^-/-^* male per cage. The male was removed after one week and the females were placed in a clean cage until delivery. F_1_ *Ggta1^+/-^* pups were weaned at 3-4 weeks of age and then co-housed until 8 weeks of age. Two or more F_1_ *Ggta1^+/-^* breeding pairs were established randomly using one male and two females per cage. F_2_ pups were weaned at 3-4 weeks of age, genotyped and segregated according to their *Ggta1^-/-^ vs. Ggta1^+/+^* genotype in separate cages until adulthood. F_3_ to F_5_ pups were generated in a similar manner. Fecal pellets from 2-3 cages per genotype were collected (10-12 weeks of age) for microbiota analysis.

The effect of *Ggta1* genotype and Ig on microbiota composition derived from *Ggta1^+/+^* mice was achieved essentially as described (Singh et al., 2020). Briefly, two or more breeding pairs were established, consisting of two *J_h_t^+/+^Ggta1^+/+^* females and one *J_h_t^-/-^Ggta1^-/-^* male per cage. The male was removed after one week and the females were placed in a clean cage until delivery. F_1_ *J_h_t^+/-^Ggta1^+/-^* pups were kept with mothers until weaning at 3-4 weeks of age and co-housed until 8 weeks of age. Two or more F_1_ *J_h_t^+/-^Ggta1^+/-^* breeding pairs were established randomly using one male and two females per cage. Littermate F_2_ pups were weaned at 3-4 weeks of age, genotyped and segregated according to their *J_h_t^+/+^Ggta1^-/-^, J_h_t^-/-^Ggta1^-/-^, J_h_t^+/+^Ggta1^+/+^* and *J_h_t^-/-^Ggta1^+/+^* genotypes in separate cages until adulthood. F_3_ pups were generated in a similar manner. Fecal pellets from 2-3 cages per genotype were collected (10-12 weeks of age) for microbiota analysis.

### Genotyping

Mice were genotyped from tail biopsies (0.5-1 cm) by PCR of genomic DNA as per manufacturer’s protocols (KAPA mouse genotyping kit #KK7352) as described (Singh et al., 2020).

### Cecal Slurry Injection

Cecal slurry injection was performed essentially as described (Singh et al., 2020). Microbiota pathogenicity experiments were performed by preparing cecal slurry from *Ggta1^-/-^ vs. Rag2^-/-^Ggta1^-/-^* mice or from *Ggta1^+/+^ vs. Rag2^-/-^Ggta^+/+^* mice in parallel and injecting to recipient mice (*i.p*. 1 mg/g body weight). Mice were monitored every 12 h for survival for 14 days or euthanized at various time points for analysis of different parameters.

### Pathogen Load

Quantification of pathogen load in the mice was performed 24 h after cecal slurry injection, essentially as described (Singh et al., 2020).

### Neutrophil depletion

Anti-Gr1 mAb (Clone: RB6-8C5, 300 μg in 200 μL PBS) was injected (i.p.) to mice 24 h before cecal slurry injection. Neutrophil depletion was confirmed by flow cytometry, using CD11b^+^Ly6G^+^ cell staining in the blood.

### Monocyte/Macrophage depletion

Clodronate or PBS liposomes (www.Clodronateliposomes.org) was injected (10 μL/g, i.p.) to mice 72 h before cecal slurry injection. Monocyte/macrophage depletion was confirmed by flow cytometry, using CD11b^+^F4/80^+^ and CD11b^+^Ly6C^+^ staining in the peritoneal lavage.

### ELISA

ELISA for IgA binding to cecal extracts was done essentially as described (Kamada et al., 2015; Zeng et al., 2016). Cecal lysate was prepared as described (Singh et al., 2020). 96-well ELISA plates (Nunc MaxiSorp #442404) were coated with the cecal lysate (100 μL/well, 4°C, overnight), washed (3x, PBS 0.05% Tween-20, Sigma-Aldrich #P7949-500ML), blocked (200 μL, PBS 1% BSA wt/vol, Calbiochem #12659-100GM, 3 h, RT) and washed (3x, PBS 0.05% Tween-20). Plates were incubated with serially diluted (50 μL) mouse sera (1:20 to 1:100 in PBS 1% BSA, wt/vol, 2 h, RT) and washed (5x, PBS/0.05% Tween-20). IgA was detected using horseradish peroxidase (HRP)-conjugated goat anti-mouse IgA (Southern Biotech #1040-05), in PBS/1 %BSA/0.01% Tween-20 (100 μL, 1:4,000 vol/vol, 1 h, RT) and plates were washed (5x, PBS/0.05% Tween-20).

For quantification of total serum and small intestinal IgA, 96-well ELISA plates (Nunc MaxiSorp #442404) were coated with goat anti-mouse IgA (Southern Biotech #1040-01, 2 μg/mL in Carbonate-Bicarbonate buffer, 100 μL/well, overnight, 4°C), washed (3x, PBS 0.05% Tween-20, Sigma-Aldrich #P7949-500ML), blocked (200 μL, PBS 1% BSA wt/vol, 2 h, RT) and washed (3x, PBS 0.05% Tween-20). Plates were incubated with serially diluted serum or gut content (50 μL, PBS 1% BSA, wt/vol, 2 h, RT) and standard mouse IgA (Southern Biotech #0106-01, prepared in duplicates) and washed (5x, PBS/0.05% Tween-20). IgA was detected using HRP-conjugated goat anti-mouse IgA (Southern Biotech #1040-05) in PBS/1%BSA/0.01% Tween-20 (100 μL, 1:4,000 vol/vol, 1 h, RT) and plates were washed (5x, PBS/0.05% Tween-20).

For quantification of anti-αGal IgG, IgM and IgA, 96-well ELISA plates (Nunc MaxiSorp #442404) were coated with αGal-BSA (Dextra, 5 μg/mL in Carbonate-Bicarbonate buffer, 50 μL/well, overnight, 4°C). Wells were washed (3x, PBS 0.05% Tween-20, Sigma-Aldrich #P7949-500ML), blocked (200 μL, PBS 1% BSA wt/vol, 2 h, RT) and washed (3x, PBS 0.05% Tween-20). Plates were incubated with serially diluted serum or fecal content (50 μL, PBS 1% BSA, wt/vol, 2 h, RT) and standard mouse anti-αGal IgG, IgM or IgA and washed (5x, PBS/0.05% Tween-20). Anti-αGal Abs were detected using HRP-conjugated goat anti-mouse IgG, IgM and IgA in PBS/1 %BSA (100 μL, 1:4000 vol/vol, 1.5 h, RT) and plates were washed (5x, PBS/0.05% Tween-20).

HRP activity was detected with 3,3’,5,5’-Tetramethylbenzidine (TMB) Substrate Reagent (BD Biosciences #555214, 50 μL, 20-25 min., dark, RT) and the reaction was stopped using sulfuric acid (2N, 50 μL). Optical densities (OD) were measured using a MultiSkan Go spectrophotometer (ThermoFisher) at λ=450 nm and normalized by subtracting background OD values (λ= 600 nm).

For measurement of fecal Lipocalin (Lcn-2), feces were resuspended with sterile PBS (100 mg/mL), vortexed (5 min.), and centrifuged (12,000 rpm, 4°C, 10 min.). Lcn-2 levels were determined in fecal supernatants using a LEGEND MAX™ Mouse NGAL (Lipocalin-2) ELISA Kit (BioLegend #443708).

### Flow cytometry of bacterial staining for IgA and αGal

Overnight cultures of bacteria were prepared as follows. Samples of 5-20 μL of each bacterial culture depending on OD_600_ measurements, and corresponding to approximately 10^6^-10^7^ cells, were fixed in PFA (4% wt/vol in PBS) and washed with filter-sterilized PBS. For detection of IgA binding and αGal expression by gut microbes, small intestinal, cecal, colon and fecal samples were homogenized (5 mg/ml in PBS) by vortexing (maximum speed, 5 min., RT) and filtered (BD Falcon™, 40 μm cell strainer # 352340). Larger debris were pelleted by centrifugation (600 g, 4°C, 5 min.). 50 μL supernatant (containing bacteria) per condition was added to a 96-well v-bottom plate (Corning Costar #3894) for staining. Bacteria were pelleted by centrifugation (3,700 g, 10 min., 4°C) and suspended in flow cytometry buffer (filter-sterilized 1xPBS, 2% BSA, wt/vol). Bacterial DNA was stained using SYTO®41 Blue Fluorescent Nucleic Acid Stain (Molecular Probes #S11352, 1:200 vol/vol, wt/vol) in flow cytometry buffer (100 μL, 30 min., RT). Cecal content from germ free (GF) mice was used as control. Bacteria were washed in flow cytometry buffer (200 μL), centrifuged (4000 g, 10 min., 4°C) and supernatant was removed by flicking the plate. Bacteria were incubated in Fluorescein Isothiocyanate (FITC)-conjugated BSI-B4 lectin from *Bandeiraea (Griffonia) simplicifolia* (Sigma-Aldrich, #L2895-1MG, 50 μL, 40 μg/mL in PBS, 2 h) for detection of αGal and anti-mouse IgA-PE mAb (mA-6E1, eBiosciences # 12-4204-82, 1:100 in PBS 2% BSA wt/vol, 30 min.) and washed as above. *E coli* O86:B7 (about 10^7^ per tube) was used as a positive control for bacterial αGal expression. Samples were re-suspended in flow cytometry buffer (300 μL), transferred to round-bottom tubes (BD Falcon™ #352235) and centrifuged (300 g, 1 min., RT). Samples were analyzed in LSR Fortessa SORP (BD Biosciences) equipped with a high-throughput sampler (HTS) using the FACSDiva Software (BD v.6.2) and analyzed by FlowJo software (v10.0.7) as described (Singh et al., 2020)..

### Extraction of bacterial DNA from feces

Bacterial DNA from fecal pellets was extracted according to manufacturer’s instructions (QIAamp Fast DNA Stool Mini Kit #50951604) as described (Singh et al., 2020).

### Amplicon Sequencing: 16S amplicons sequencing and analysis

The 16S rRNA V4 region was amplified and sequenced following the Earth Microbiome Project (http://www.earthmicrobiome.org/emp-standard-protocols/), and analyzed using QIIME 1.9.1 as described (Singh et al., 2020). 16S sequencing data was submitted to the Sequence Read Archive (SRA), with the BioProject reference PRJNA701192.

### Statistical analysis

All statistical tests were performed using GraphPad Prism Software (v.6.0). All statistical details of experiments including statistical tests, exact value of n, what n represents, definition of center, dispersion and precision measures are provided in each figure legend.

## Data availability

All data generated during this study are included in the manuscript and supporting files. 16S sequencing data was submitted to the Sequence Read Archive (SRA), with the BioProject reference PRJNA701192.

## Acknowledgements

We thank our colleagues J. Howard and I. Gordo (Instituto Gulbenkian de Ciência; IGC) for critical review of the manuscript, IGC Genomics, Flow Cytometry, Antibody, Histopathology and Animal House Facilities. S.S. was supported by Fundac□ão para a Cie□ncia e Tecnologia (FCT; SFRH/BD/52177/2013), J.A.T. by an ESCMID Research Grant and FCT (SFRH/BPD/112135/2015) and M.P.S. by the Gulbenkian, “La Caixa” (HR18-00502) and Bill & Melinda Gates (OPP1148170) Foundations and FCT (5723/2014 and FEDER029411).

## Author contributions

S.S. contributed critically to study design and performed most experimental work and data analyses. P.B.A. performed flow cytometry analysis of probiotic bacterial strains with B.Y., and experiments pertaining to the role of IgA *in vivo*. J.A.T. performed flow cytometry analysis of mouse microbiota and contributed to study interpretation. S.C. generated mouse strains with S.S. D.S. and M.T. performed 16S rRNA sequencing analysis. M.P.S. drove the study design and wrote the manuscript with S.S.

## Supplementary Figure legends

**Suppl. Figure 1. Analyses of gut microbiota composition of *Ggta1^+/+^* and *Ggta1^-/-^* mice.** Relative abundance of bacteria at all levels of taxonomy, present at >2% frequency, in the same mice as in Figure 1A. Symbols represent individual mice. Red bars correspond to mean values. Error bars correspond to SD. Adjusted P values calculated using Benjamini-Hochberg correction. *P < 0.05, **P < 0.01, ***P < 0.005, ****P < 0.001.

**Suppl. Figure 2. Analyses of gut microbiota composition of *Ggta1^+/+^ vs. Ggta1^-/-^* mice.** Relative abundance of bacteria at all levels of taxonomy, present at >2% frequency in the same mice as in Figure 1A. Stacked bars represent the mean of the bacterial taxa. Colors represent the relative fraction of each taxon. Data from 1 experiment.

**Suppl. Figure 3. *Ggta1* deletion does not cause inflammation at steady state. A**) Representative H/E sections of the small intestine, large intestine, liver, spleen, lung and kidney of *Ggta1^+/+^* (n=5) and *Ggta1^-/-^* (n=5) mice at steady state. **B**) Fecal Lcn-2 concentrations in *Ggta1^+/+^* (n=10) and *Ggta1^-/-^* (n=10) mice at steady state. Symbols represent individual mice. Red bars correspond to mean values. Error bars correspond to SD, ns: not significant.

**Suppl. Figure 4. *Ggta1* deletion shapes the gut microbiota composition.** Relative abundance of bacteria at all levels of taxonomy, present at >2% frequency in the same mice as in Figure 1C-F. Stacked bars represent the mean of the bacterial taxa. Colors represent the relative fraction of each taxon. Data from 1 experiment.

**Suppl. Figure 5. *Ggta1* deletion enhances IgA responses to the gut microbiota. A**) Concentration of total IgA in serum of *Ggta1^+/+^* (n=9), GF *Ggta1^+/+^* (n=5), *Ggta1^-/-^* (n=5) and GF *Ggta1^-/-^* (n=6) mice. **B**) Concentration of total IgA in the small intestinal content of *Ggta1^+/+^* (n=10), GF *Ggta1^+/+^* (n=5), *Ggta1^-/-^* (n=10) and GF *Ggta1^-/-^* (n=5) mice. **C**) Relative binding of IgA in the serum of GF *Ggta1^+/+^* (n=5) and GF *Ggta1^-/-^* (n=5) mice to fecal extract from *Ggta1^+/+^* mice at indicated time-points after colonization with cecal extract from *Ggta1^+/+^* mice; 2 experiments. **D**) Concentration of total IgA in serum of *Ggta1^-/-^* (n=7), *Tcrβ^-/-^ Ggta1^-/-^* (n=12) mice. **E**) Representative plots showing the gating strategy for staining of intestinal content with Syto41, BSI-B4 and Anti-IgA. **F**) Quantification of αGal^+^, IgA^+^ and IgA^+^αGal^+^ bacteria in the cecum, colon and feces of the same mice as in Figure 2D-E. Symbols (A, B, C, D, F) are individual mice. Red bars (A, B, C, D, F) correspond to mean values. Error bars (A, B, C, D, F) correspond to SD. P values in (A, B, C) calculated using Kruskal-Wallis test using Dunn’s multiple comparisons test and in (D, F) using Mann-Whitney test. *P < 0.05, **P < 0.01, ns: not significant.

**Suppl. Figure 6. αGal expression by probiotic bacteria.** Representative flow cytometry plots of the data in Figure 2C showing probiotic bacterial strains unstained or stained with BSI-B4 lectin.

**Suppl. Figure 7. *Ggta1* deletion shapes the gut microbiota via an Ig-dependent mechanism.** Microbiota Principal Coordinate Analysis of **A**) Weighted and Unweighted Unifrac and **B**) Distance of Weighted and Unweighted Unifrac of 16S rRNA amplicons, in fecal samples from F_2_ *J_h_t^+/+^Ggta1^+/+^* (n=8) *vs. J_h_t^-/-^Ggta1^+/+^* (n=11) mice and F_2_ *J_h_t^+/+^Ggta1^-/-^* (n=5) *vs. J_h_t^-/-^Ggta1^-/-^* (n=8) mice generated as described in Figure 3A. **C-D**) Cladogram and linear discriminant analysis (LDA) scores generated from LEfSe analysis, representing taxa enriched in the fecal microbiota of the same **C**) F_3_ *J_h_t^+/+^Ggta1^+/+^* vs*. J_h_t^+/+^Ggta1^-/-^* mice and **D**) F_3_ *J_h_t^-/-^Ggta1^+/+^ vs. J_h_t^+/+^Ggta1^+/+^* mice as in Figure 3B-D. Symbols (A, B) are individual mice. Red bars (B) correspond to mean values. Error bars (B) correspond to SD. P values in (A) calculated using PERMANOVA and in (B) using Mann-Whitney test. ns: not significant.

**Suppl. Figure 8. *Ggta1* deletion reduces microbiota pathogenicity. A**) Schematic showing infection of *Ggta1^-/-^* mice with a cecal inoculum from either *Ggta1^-/-^* mice or *Rag2^-/-^Ggta1^-/-^* mice. **B**) Survival of *Ggta1^-/-^* (n=14) mice after infection with a cecal inoculum from *Ggta1^-/-^* mice and of *Ggta1^-/-^* (n=13) mice after infection with a cecal inoculum from *Rag2^-/-^Ggta1^-/-^* mice; 2 experiments. **C**) Survival of *J_h_t^+/+^Ggta1^-/-^* (n=9) and *J_h_t^-/-^Ggta1^-/-^* (n=8) mice after infection with a cecal inoculum from *Ggta1^-/-^* mice; 2 experiments. **D**) Survival of *Tcrβ^+/+^Ggta1^-/-^* (n=8) and *Tcrβ^+/+^ Ggta1^-/-^* (n=12) mice infected as in (C); 2 experiments. **E**) Survival of *Rag2^+/+^Ggta1^-/-^* (n=9) and *Rag2^-/-^ Ggta1^-/-^* (n=23) mice infected as in (C); 5 experiments. **F**) Representative plots showing depletion of CD11b^+^Ly6G^+^ cells in the blood 24 hours after Anti-Gr1 Ab injection (i.p.) in the same mice as in Figure 4G. **G**) Representative plots showing depletion of CD11b^+^F4/80^+^ and CD11b^+^Ly6C^+^ cells in the peritoneal lavage 72 hours after injection (i.p.) with Clodronate liposomes. **H**) Survival of *Ggta1^-/-^* mice receiving PBS liposomes (n=7) or Clodronate liposomes (n=8), 72 hours before infection (i.p.) with cecal inoculum from *Ggta1^-/-^* mice; 2 experiments. P values in (B, C, D, E, H) calculated with log-rank test. Peritoneal cavity (PC). ns: not significant.

## References

Allison, A.C. (1954). Protection afforded by sickle-cell trait against subtertian malareal infection. Br Med J 1, 290–294.

Atherton, J.C., and Blaser, M.J. (2009). Coadaptation of Helicobacter pylori and humans: ancient history, modern implications. The Journal of clinical investigation 119, 2475–2487.

Ayres, J.S. (2016). Cooperative Microbial Tolerance Behaviors in Host-Microbiota Mutualism. Cell 165, 1323–1331.

Bernth Jensen, J.M., Skeldal, S., Petersen, M.S., Møller, B.K., Hoffmann, S., Jensenius, J.C., Skov Sørensen, U.B., and Thiel, S. (2020). The human natural anti-αGal antibody targets common pathogens by broad-spectrum polyreactivity. Immunology n/a.

Buffie, C.G., and Pamer, E.G. (2013). Microbiota-mediated colonization resistance against intestinal pathogens. Nat Rev Immunol 13, 790–801.

Bunker, J.J., Erickson, S.A., Flynn, T.M., Henry, C., Koval, J.C., Meisel, M., Jabri, B., Antonopoulos, D.A., Wilson, P.C., and Bendelac, A. (2017). Natural polyreactive IgA antibodies coat the intestinal microbiota. Science 358.

Bunker, J.J., Flynn, T.M., Koval, J.C., Shaw, D.G., Meisel, M., McDonald, B.D., Ishizuka, I.E., Dent, A.L., Wilson, P.C., Jabri, B., et al. (2015). Innate and Adaptive Humoral Responses Coat Distinct Commensal Bacteria with Immunoglobulin A. Immunity 43, 541–553.

Caruso, R., Lo, B.C., and Núñez, G. (2020). Host–microbiota interactions in inflammatory bowel disease. Nature Reviews Immunology 20, 411–426.

Chassaing, B., Srinivasan, G., Delgado, M.A., Young, A.N., Gewirtz, A.T., and Vijay-Kumar, M. (2012). Fecal lipocalin 2, a sensitive and broadly dynamic non-invasive biomarker for intestinal inflammation. PLoS One 7, e44328.

Chow, J., Tang, H., and Mazmanian, S.K. (2011). Pathobionts of the gastrointestinal microbiota and inflammatory disease. Curr Opin Immunol 23, 473–480.

Daley, J.M., Thomay, A.A., Connolly, M.D., Reichner, J.S., and Albina, J.E. (2008). Use of Ly6G-specific monoclonal antibody to deplete neutrophils in mice. Journal of leukocyte biology 83, 64–70.

Dethlefsen, L., McFall-Ngai, M., and Relman, D.A. (2007). An ecological and evolutionary perspective on human–microbe mutualism and disease. Nature 449, 811–818.

Donaldson, G.P., Ladinsky, M.S., Yu, K.B., Sanders, J.G., Yoo, B.B., Chou, W.-C., Conner, M.E., Earl, A.M., Knight, R., Bjorkman, P.J., et al. (2018). Gut microbiota utilize immunoglobulin A for mucosal colonization. Science 360, 795–800.

Fagarasan, S., Kawamoto, S., Kanagawa, O., and Suzuki, K. (2010). Adaptive immune regulation in the gut: T cell-dependent and T cell-independent IgA synthesis. Annual review of immunology 28, 243–273.

Galili, U. (2016). Natural anticarbohydrate antibodies contributing to evolutionary survival of primates in viral epidemics? Glycobiology In press.

Galili, U. (2019). Evolution in primates by “Catastrophic-selection” interplay between enveloped virus epidemics, mutated genes of enzymes synthesizing carbohydrate antigens, and natural anti-carbohydrate antibodies. Am J Phys Anthropol 168, 352–363.

Galili, U., Clark, M.R., Shohet, S.B., Buehler, J., and Macher, B.A. (1987). Evolutionary relationship between the natural anti-Gal antibody and the Gal alpha 1----3Gal epitope in primates. Proc Natl Acad Sci U S A 84, 1369–1373.

Galili, U., Shohet, S.B., Kobrin, E., Stults, C.L., and Macher, B.A. (1988). Man, apes, and Old World monkeys differ from other mammals in the expression of alpha-galactosyl epitopes on nucleated cells. J Biol Chem 263, 17755–17762.

Gálvez, E.J.C., Iljazovic, A., Gronow, A., Flavell, R., and Strowig, T. (2017). Shaping of Intestinal Microbiota in Nlrp6- and Rag2-Deficient Mice Depends on Community Structure. Cell Rep 21, 3914–3926.

Gensollen, T., Iyer, S.S., Kasper, D.L., and Blumberg, R.S. (2016). How colonization by microbiota in early life shapes the immune system. Science 352, 539–544.

Ghaderi, D., Springer, S.A., Ma, F., Cohen, M., Secrest, P., Taylor, R.E., Varki, A., and Gagneux, P. (2011). Sexual selection by female immunity against paternal antigens can fix loss of function alleles. Proc Natl Acad Sci U S A 108, 17743–17748.

Gomez de Agüero, M., Ganal-Vonarburg, S.C., Fuhrer, T., Rupp, S., Uchimura, Y., Li, H., Steinert, A., Heikenwalder, M., Hapfelmeier, S., Sauer, U., et al. (2016). The maternal microbiota drives early postnatal innate immune development. Science 351, 1296–1302.

Gu, H., Zou, Y.R., and Rajewsky, K. (1993). Independent control of immunoglobulin switch recombination at individual switch regions evidenced through Cre-loxP-mediated gene targeting. Cell 73, 1155–1164.

Guo, H., Yi, W., Shao, J., Lu, Y., Zhang, W., Song, J., and Wang, P.G. (2005). Molecular analysis of the O-antigen gene cluster of Escherichia coli O86:B7 and characterization of the chain length determinant gene (wzz). Appl Environ Microbiol 71, 7995–8001.

Haak, B.W., and Wiersinga, W.J. (2017). The role of the gut microbiota in sepsis. The lancet Gastroenterology & hepatology 2, 135–143.

Haldane, J.B.S. (1949). Disease and evolution, Ricercha.

Hamadeh, R.M., Estabrook, M.M., Zhou, P., Jarvis, G.A., and Griffiss, J.M. (1995a). Anti-Gal binds to pili of Neisseria meningitidis: the immunoglobulin A isotype blocks complement-mediated killing. Infection and immunity 63, 4900–4906.

Hamadeh, R.M., Galili, U., Zhou, P., and Griffiss, J.M. (1995b). Anti-alpha-galactosyl immunoglobulin A (IgA), IgG, and IgM in human secretions. Clin Diagn Lab Immunol 2, 125–131.

Hooper, L.V., Littman, D.R., and Macpherson, A.J. (2012). Interactions Between the Microbiota and the Immune System. Science 336, 1268–1273.

Huttenhower, C., Gevers, D., Knight, R., Abubucker, S., Badger, J.H., Chinwalla, A.T., Creasy, H.H., Earl, A.M., FitzGerald, M.G., Fulton, R.S., et al. (2012). Structure, function and diversity of the healthy human microbiome. Nature 486, 207–214.

Kamada, N., Sakamoto, K., Seo, S.U., Zeng, M.Y., Kim, Y.G., Cascalho, M., Vallance, B.A., Puente, J.L., and Nunez, G. (2015). Humoral Immunity in the Gut Selectively Targets Phenotypically Virulent Attaching-and-Effacing Bacteria for Intraluminal Elimination. Cell Host Microbe 17, 617–627.

Kawamoto, S., Maruya, M., Kato, Lucia M., Suda, W., Atarashi, K., Doi, Y., Tsutsui, Y., Qin, H., Honda, K., Okada, T., et al. (2014). Foxp3+ T Cells Regulate Immunoglobulin A Selection and Facilitate Diversification of Bacterial Species Responsible for Immune Homeostasis. Immunity 41, 152–165.

Koch, M.A., Reiner, G.L., Lugo, K.A., Kreuk, L.S., Stanbery, A.G., Ansaldo, E., Seher, T.D., Ludington, W.B., and Barton, G.M. (2016). Maternal IgG and IgA Antibodies Dampen Mucosal T Helper Cell Responses in Early Life. Cell 165, 827–841.

Kubinak, J.L., and Round, J.L. (2016). Do antibodies select a healthy microbiota? Nat Rev Immunol 16, 767–774.

Lane-Petter, W. (1962). The Provision and Use of Pathogen-Free Laboratory Animals. Proceedings of the Royal Society of Medicine 55, 253–263.

Lewis, H. (1962). Catastrophic Selection as a Factor in Speciation. Evolution 16, 257–271.

Linz, B., Balloux, F., Moodley, Y., Manica, A., Liu, H., Roumagnac, P., Falush, D., Stamer, C., Prugnolle, F., van der Merwe, S.W., et al. (2007). An African origin for the intimate association between humans and Helicobacter pylori. Nature 445, 915–918.

Lozupone, C., and Knight, R. (2005). UniFrac: a New Phylogenetic Method for Comparing Microbial Communities. Applied and Environmental Microbiology 71, 8228–8235.

Macpherson, A.J., Gatto, D., Sainsbury, E., Harriman, G.R., Hengartner, H., and Zinkernagel, R.M. (2000). A primitive T cell-independent mechanism of intestinal mucosal IgA responses to commensal bacteria. Science 288, 2222–2226.

Macpherson, A.J., Yilmaz, B., Limenitakis, J.P., and Ganal-Vonarburg, S.C. (2018). IgA Function in Relation to the Intestinal Microbiota. Annual review of immunology 36, 359–381.

McFall-Ngai, M. (2007). Care for the community. Nature 445, 153–153.

McLoughlin, K., Schluter, J., Rakoff-Nahoum, S., Smith, Adrian L., and Foster, Kevin R. (2016). Host Selection of Microbiota via Differential Adhesion. Cell Host & Microbe 19, 550–559.

Moeller, A.H., Caro-Quintero, A., Mjungu, D., Georgiev, A.V., Lonsdorf, E.V., Muller, M.N., Pusey, A.E., Peeters, M., Hahn, B.H., and Ochman, H. (2016). Cospeciation of gut microbiota with hominids. Science 353, 380–382.

Montassier, E., Al-Ghalith, G.A., Mathé, C., Le Bastard, Q., Douillard, V., Garnier, A., Guimon, R., Raimondeau, B., Touchefeu, Y., Duchalais, E., et al. (2019). Distribution of Bacterial α1,3-Galactosyltransferase Genes in the Human Gut Microbiome. Front Immunol 10, 3000.

Moor, K., Diard, M., Sellin, M.E., Felmy, B., Wotzka, S.Y., Toska, A., Bakkeren, E., Arnoldini, M., Bansept, F., Co, A.D., et al. (2017). High-avidity IgA protects the intestine by enchaining growing bacteria. Nature 544, 498–502.

Palm, N.W., de Zoete, M.R., Cullen, T.W., Barry, N.A., Stefanowski, J., Hao, L., Degnan, P.H., Hu, J., Peter, I., Zhang, W., et al. (2014). Immunoglobulin A coating identifies colitogenic bacteria in inflammatory bowel disease. Cell 158, 1000–1010.

Peterson, D.A., McNulty, N.P., Guruge, J.L., and Gordon, J.I. (2007). IgA Response to Symbiotic Bacteria as a Mediator of Gut Homeostasis. Cell Host & Microbe 2, 328–339.

Pickard, J.M., Maurice, C.F., Kinnebrew, M.A., Abt, M.C., Schenten, D., Golovkina, T.V., Bogatyrev, S.R., Ismagilov, R.F., Pamer, E.G., Turnbaugh, P.J., et al. (2014). Rapid fucosylation of intestinal epithelium sustains host-commensal symbiosis in sickness. Nature 514, 638–641.

Repik, P.M., Strizki, J.M., and Galili, U. (1994). Differential host-dependent expression of alpha-galactosyl epitopes on viral glycoproteins: a study of eastern equine encephalitis virus as a model. J Gen Virol 75 (Pt 5), 1177–1181.

Rudd, K.E., Johnson, S.C., Agesa, K.M., Shackelford, K.A., Tsoi, D., Kievlan, D.R., Colombara, D.V., Ikuta, K.S., Kissoon, N., Finfer, S., et al. (2020). Global, regional, and national sepsis incidence and mortality, 1990-2017: analysis for the Global Burden of Disease Study. Lancet 395, 200–211.

Singer, M., Deutschman, C.S., Seymour, C.W., Shankar-Hari, M., Annane, D., Bauer, M., Bellomo, R., Bernard, G.R., Chiche, J.D., Coopersmith, C.M., et al. (2016). The Third International Consensus Definitions for Sepsis and Septic Shock (Sepsis-3). JAMA 315, 801–810.

Singh, S., Thompson, J.A., Weis, S., Sobral, D., Truglio, M., Yilmaz, B., Rebelo, S., Cardoso, S., Gjini, E., Nuñez, G., et al. (2020). A trade-off between resistance to infection and reproduction in primate evolution. bioRxiv, 2020.2007.2010.186742.

Soares, M.P., and Yilmaz, B. (2016). Microbiota Control of Malaria Transmission. Trends Parasitol 32, 120–130.

Sonnenburg, E.D., Smits, S.A., Tikhonov, M., Higginbottom, S.K., Wingreen, N.S., and Sonnenburg, J.L. (2016). Diet-induced extinctions in the gut microbiota compound over generations. Nature 529, 212–215.

Spor, A., Koren, O., and Ley, R. (2011). Unravelling the effects of the environment and host genotype on the gut microbiome. Nature Reviews Microbiology 9, 279–290.

Springer, G.F., and Horton, R.E. (1969). Blood group isoantibody stimulation in man by feeding blood group-active bacteria. J Clin Invest 48, 1280–1291.

Stearns, S.C., and Medzhitov, R. (2015). Evolutionary Medicine, 1st Edition edn (Oxford University press).

Sunderkötter, C., Nikolic, T., Dillon, M.J., Van Rooijen, N., Stehling, M., Drevets, D.A., and Leenen, P.J. (2004a). Subpopulations of mouse blood monocytes differ in maturation stage and inflammatory response. J Immunol 172, 4410–4417.

Sunderkötter, C., Nikolic, T., Dillon, M.J., van Rooijen, N., Stehling, M., Drevets, D.A., and Leenen, P.J.M. (2004b). Subpopulations of Mouse Blood Monocytes Differ in Maturation Stage and Inflammatory Response. The Journal of Immunology 172, 4410–4417.

Takeuchi, Y., Porter, C.D., Strahan, K.M., Preece, A.F., Gustafsson, K., Cosset, F.L., Weiss, R.A., and Collins, M.K. (1996). Sensitization of cells and retroviruses to human serum by (alpha 1-3) galactosyltransferase. Nature 379, 85–88.

Tearle, R.G., Tange, M.J., Zannettino, Z.L., Katerelos, M., Shinkel, T.A., Van Denderen, B.J., Lonie, A.J., Lyons, I., Nottle, M.B., Cox, T., et al. (1996). The alpha-1,3-galactosyltransferase knockout mouse. Implications for xenotransplantation. Transplantation 61, 13–19.

Ubeda, C., Lipuma, L., Gobourne, A., Viale, A., Leiner, I., Equinda, M., Khanin, R., and Pamer, E.G. (2012). Familial transmission rather than defective innate immunity shapes the distinct intestinal microbiota of TLR-deficient mice. J Exp Med 209, 1445–1456.

Vincent, J.L., Rello, J., Marshall, J., Silva, E., Anzueto, A., Martin, C.D., Moreno, R., Lipman, J., Gomersall, C., Sakr, Y., et al. (2009). International study of the prevalence and outcomes of infection in intensive care units. JAMA 302, 2323–2329.

Vonaesch, P., Anderson, M., and Sansonetti, P.J. (2018). Pathogens, microbiome and the host: emergence of the ecological Koch’s postulates. FEMS microbiology reviews 42, 273–292.

Yilmaz, B., Portugal, S., Tran, T.M., Gozzelino, R., Ramos, S., Gomes, J., Regalado, A., Cowan, P.J., d’Apice, A.J., Chong, A.S., et al. (2014). Gut Microbiota Elicits a Protective Immune Response against Malaria Transmission. Cell 159, 1277–1289.

Zeng, Melody Y., Cisalpino, D., Varadarajan, S., Hellman, J., Warren, H.S., Cascalho, M., Inohara, N., and Núñez, G. (2016). Gut Microbiota-Induced Immunoglobulin G Controls Systemic Infection by Symbiotic Bacteria and Pathogens. Immunity 44, 647–658.

