## Supplementary figures and images for "Glycan-Based Shaping Of The Microbiota During Primate Evolution"

### Supplementary Figure 1

Figure S1 Singh et al.

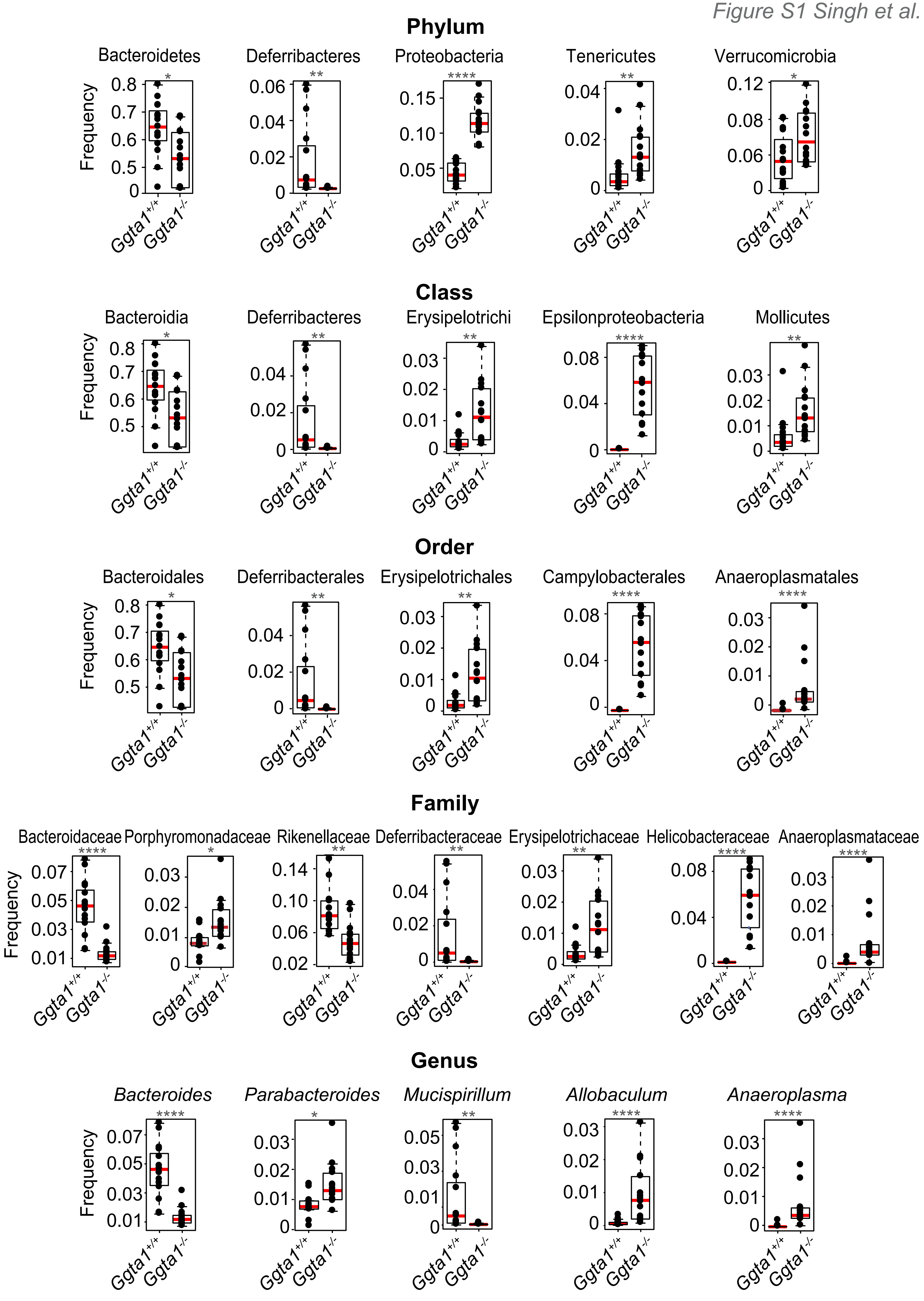

### Supplementary Figure 2

*Figure S2 Singh et al.*

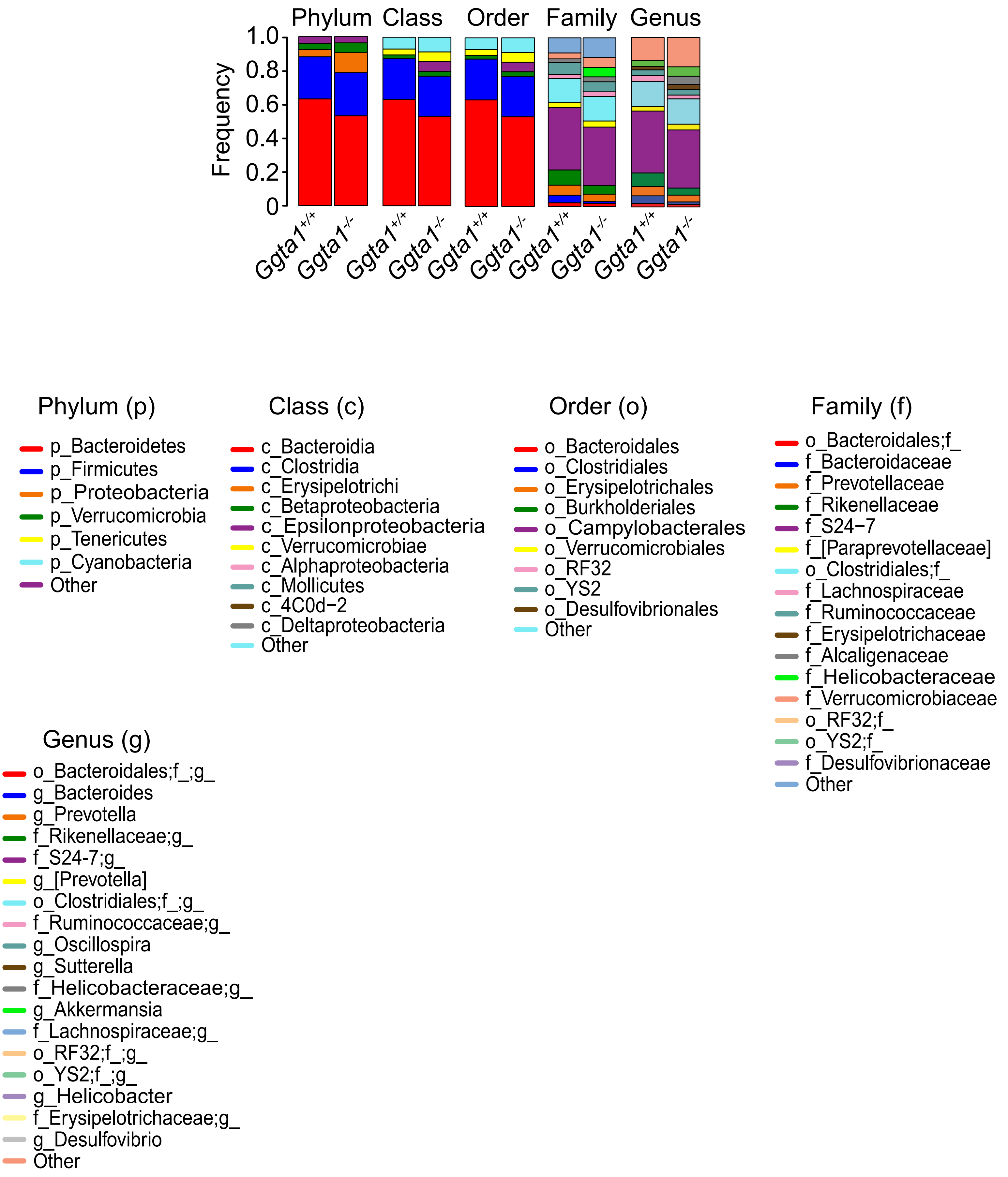

### Supplementary Figure 3

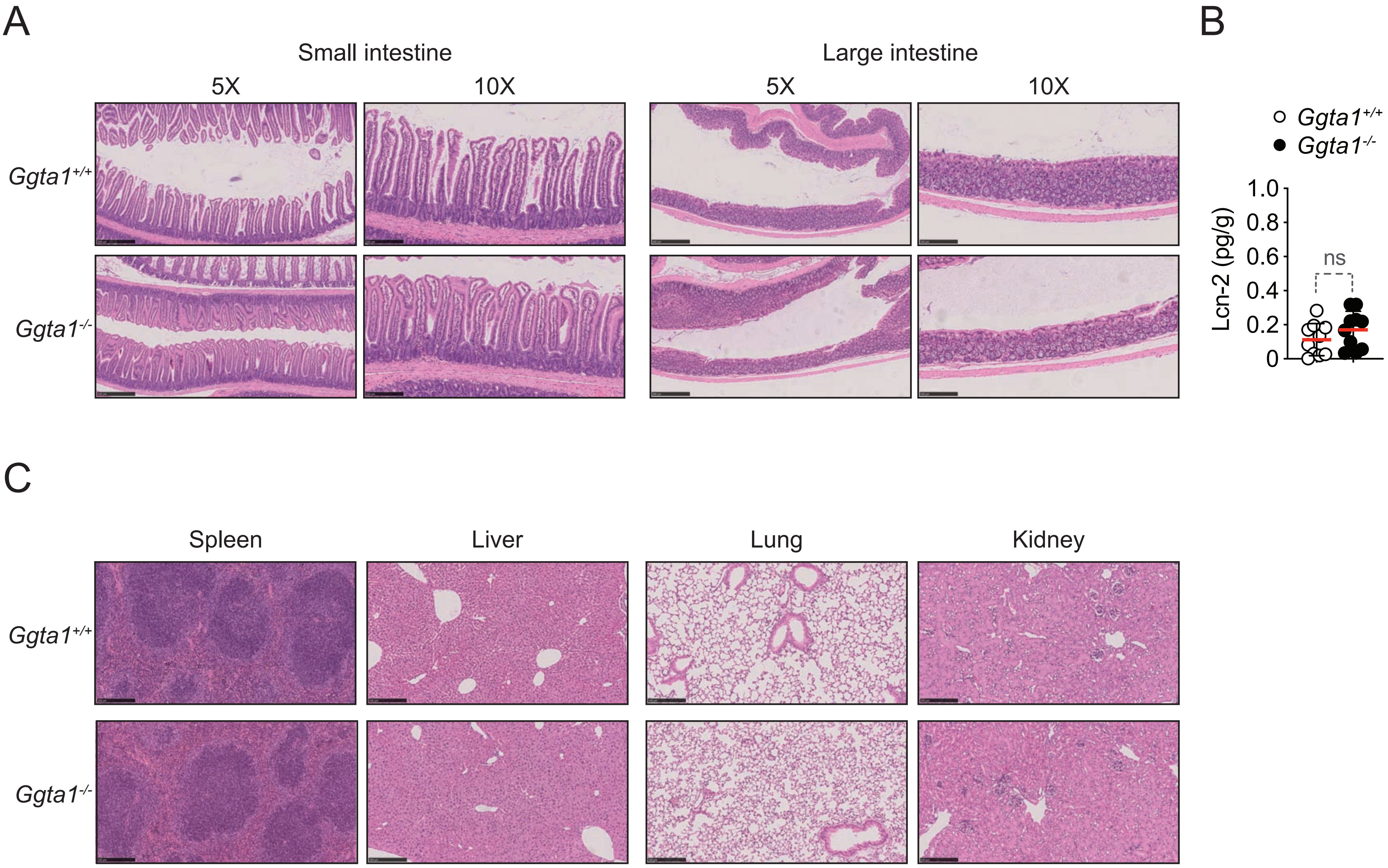

### Supplementary Figure 4

*Figure S4 Singh et al.*

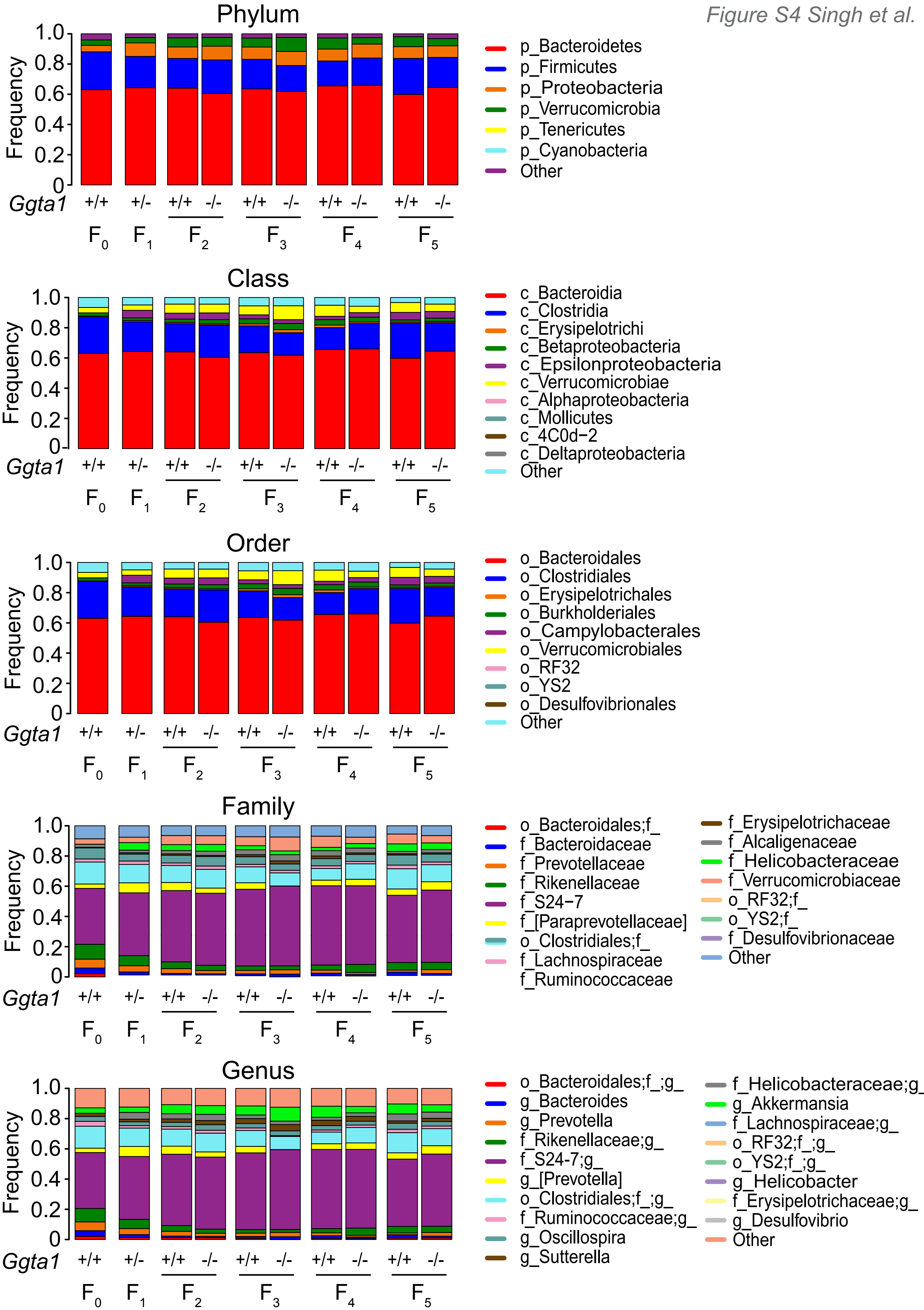

### Supplementary Figure 5

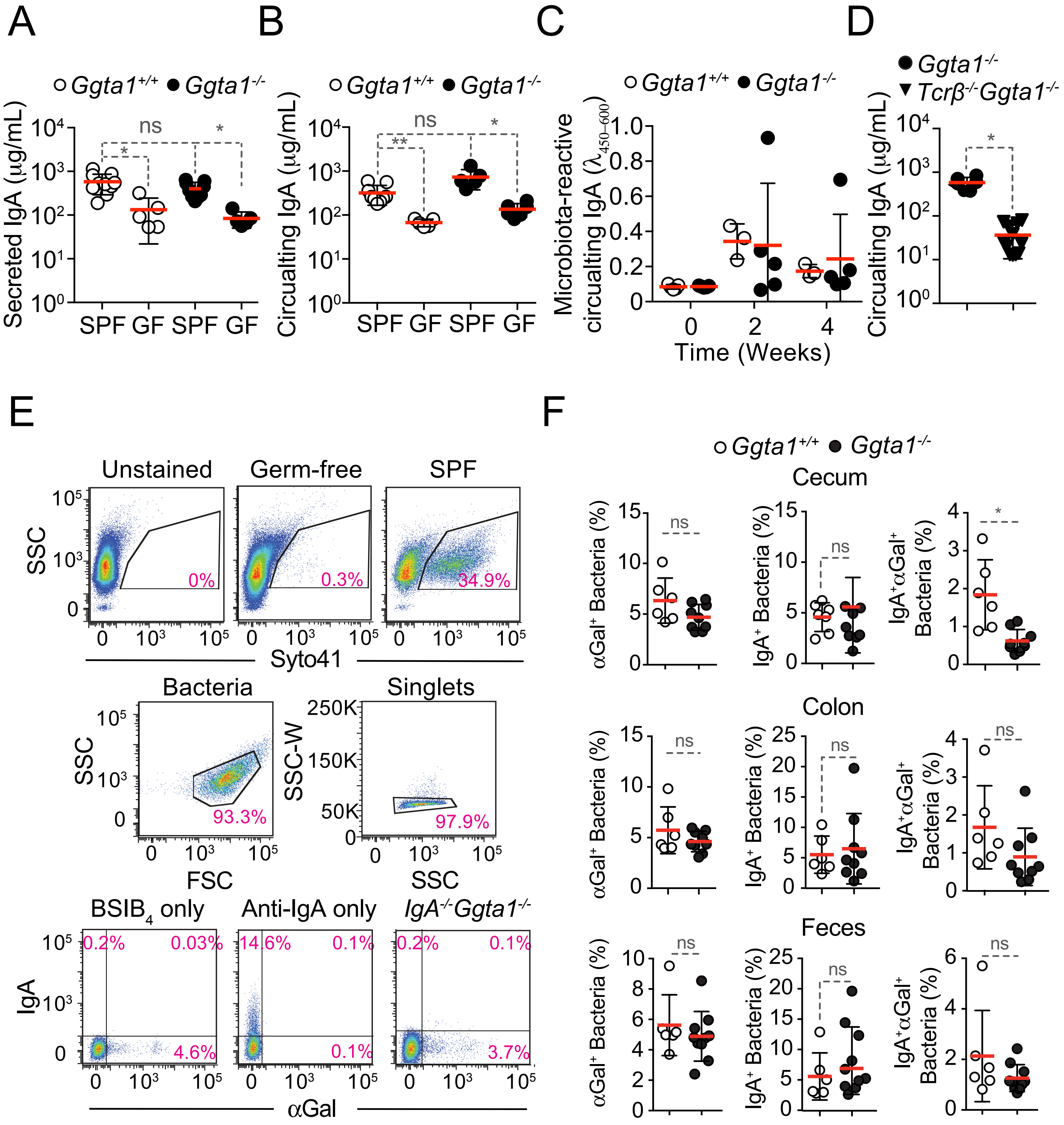

### Supplementary Figure 6

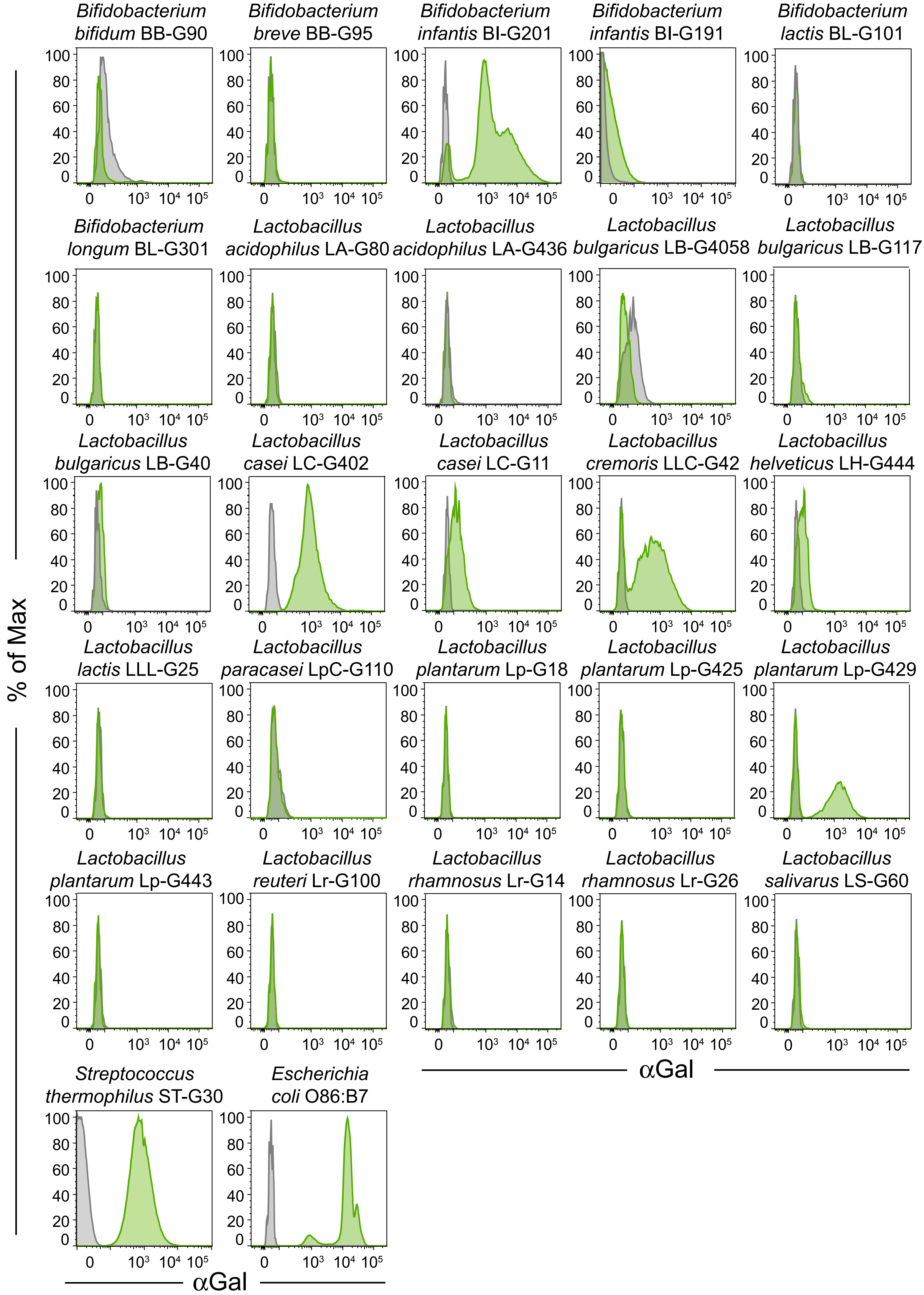

### Supplementary Figure 7

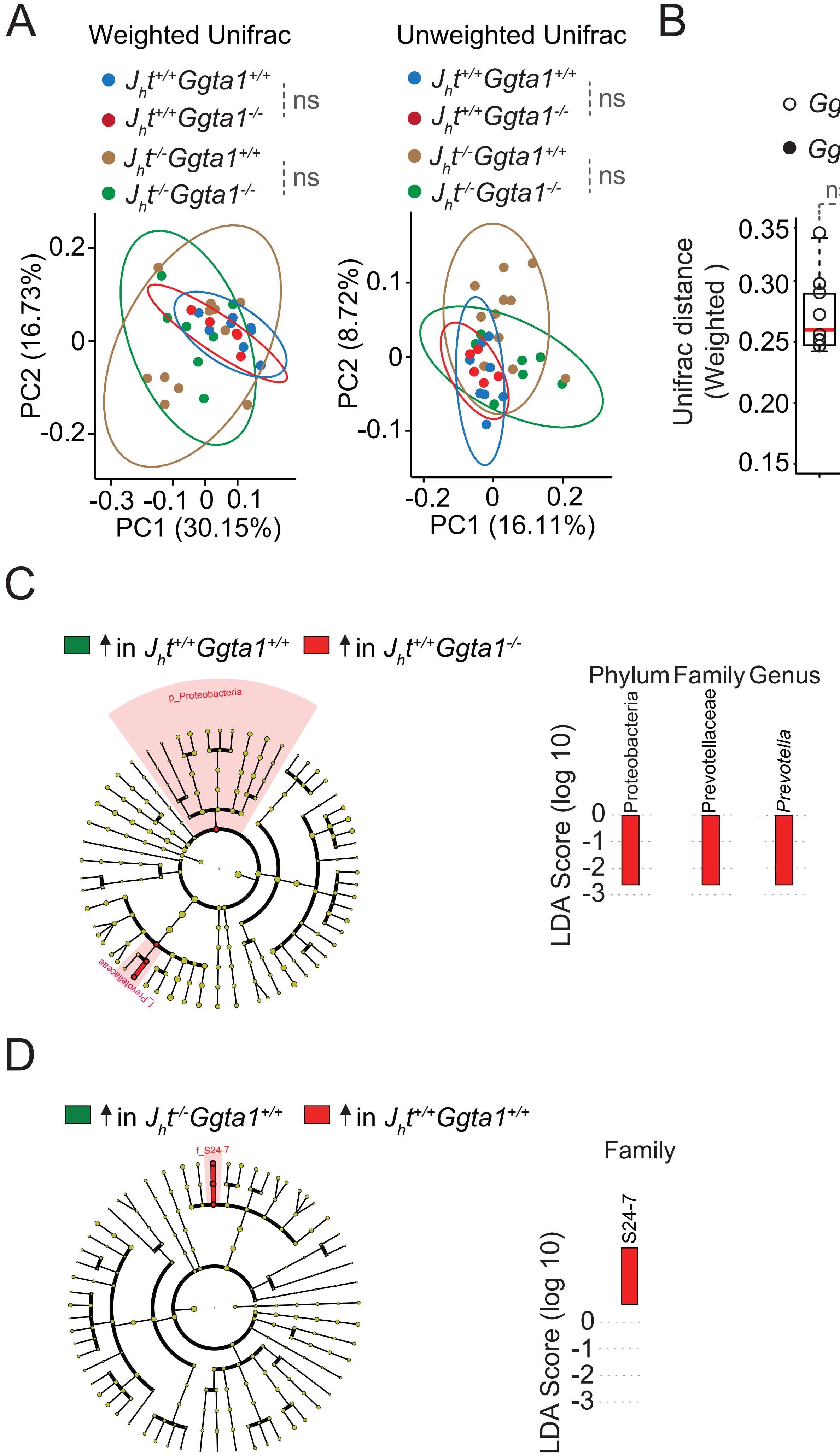

### Supplementary Figure 8

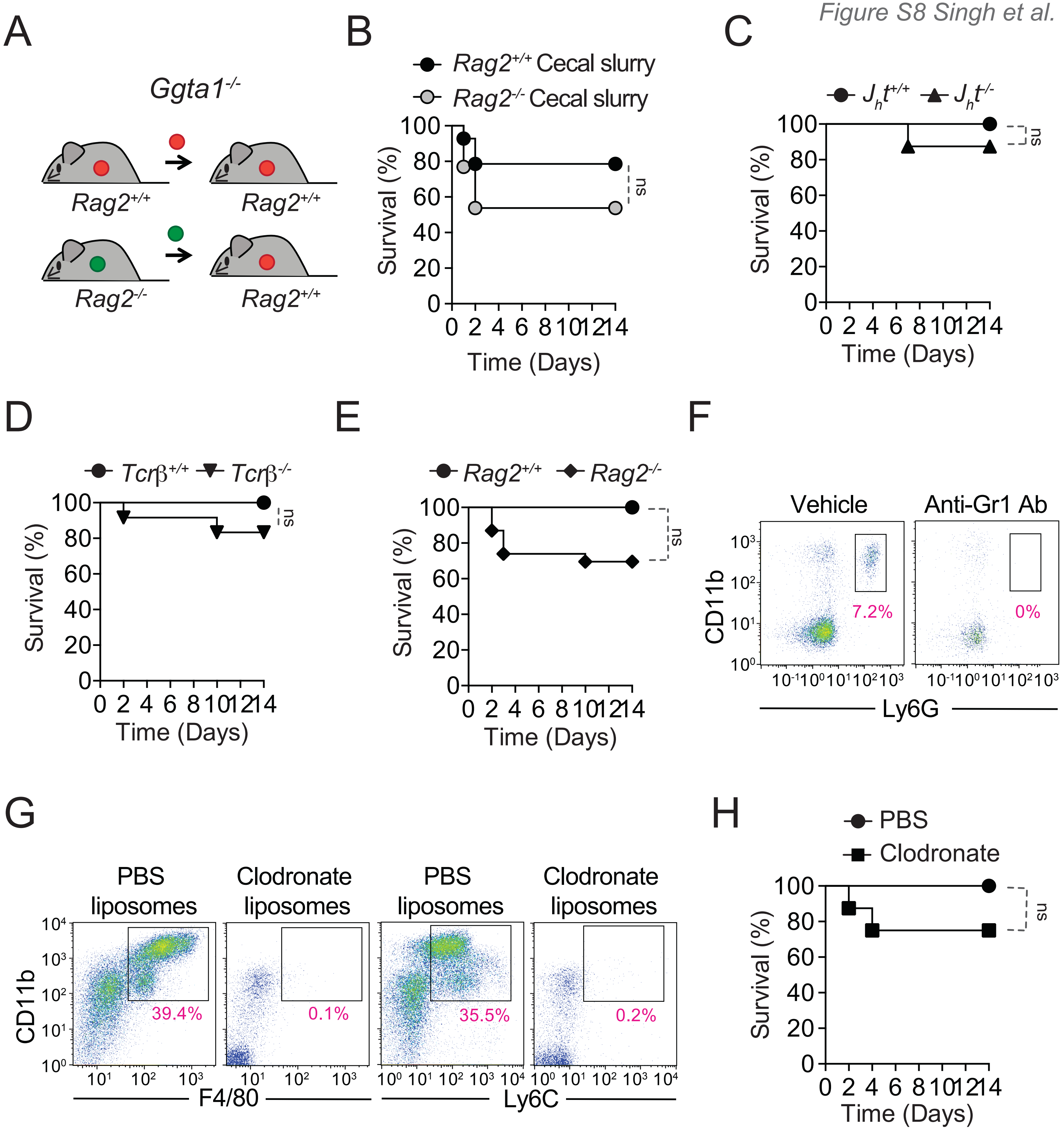
